# Screening for functional circular RNAs using the CRISPR-Cas13 system

**DOI:** 10.1101/2020.03.23.002865

**Authors:** Siqi Li, Xiang Li, Wei Xue, Lin Zhang, Shi-Meng Cao, Yun-Ni Lei, Liang-Zhong Yang, Si-Kun Guo, Jia-Lin Zhang, Xiang Gao, Jia Wei, Jinsong Li, Li Yang, Ling-Ling Chen

**Author notes:** These authors contributed equally (Siqi Li, Xiang Li, Wei Xue, Lin Zhang). Correspondence should be addressed to L.Y., J. L. and L.-L.C.

## Abstract

Circular RNAs (circRNAs) produced from back-spliced exons are widely expressed, but individual circRNA functions remain poorly understood due to inadequate methods, such as RNAi and genome engineering, in distinguishing overlapped exons in circRNAs from those in linear cognate mRNAs^1,2^. Here we report that the programable RNA-guided, RNA-targeting CRISPR-Cas13, RfxCas13d, effectively and specifically discriminates circRNAs from mRNAs, using guide (g)RNAs targeting sequences spanning the back-splicing junction (BSJ) sites featured in RNA circles. Using a lentiviral library that targets sequences across BSJ sites of highly expressed human circRNAs, we show that a group of circRNAs are important for cell growth mostly in a cell-type specific manner and that a common oncogenic circRNA, *circFAM120A,* promotes cell proliferation *in vitro* and *in vivo* by preventing *FAM120A* mRNA from binding the translation inhibitor IGF2BP2 for efficient translation. Application of RfxCas13d/BSJ-gRNA screening has also uncovered *circMan1a2* with regulatory potential in mouse embryo preimplantation development. Together, these results establish CRISPR-RfxCas13d as a useful tool for the discovery and functional study of circRNAs at both individual and large-scale levels.

## Introduction

Back-splicing (BS) of pre-mRNA exons leads to genome-wide expression of circRNAs that are identical to linear mRNA sequences except at the BS junction (BSJ) sites. Their functions largely remain elusive, partly owing to the lack of effective tools to interfere with circular but not linear RNA levels in cells^1^. Short hairpin RNAs (shRNAs) or specific small interfering RNAs (siRNAs) targeting BSJ sites of circRNAs have been used to knock down circRNAs^3–5^. However, even partial complementarity of a half-RNAi (~10 nt) to linear mRNAs may affect parental gene expression (Extended Data Fig. 1a). Genome engineering has been used to deplete a circRNA by removing either the entire circRNA-forming exon^6^ or the intronic *cis*-elements required for circRNA biogenesis^7,8^ (Extended Data Fig. 1b), but only a few circRNAs were studied with these approaches^6–8^. A simple and scalable tool that can be used to identify functional circRNAs is still lacking.

Recent studies have established that RNA-targeting type VI CRISPR effectors, known as Cas13a, Cas13b and Cas13d RNases^9–12^, can be directed to cleave single-stranded (ss) RNA targets carrying complementary protospacers by a CRISPR RNA in mammalian cells^11–14^. Type VI CRISPR-Cas knockdown requires 22- to 30-nt spacers (guide RNAs, gRNAs), exhibits higher specificity than RNAi, and is intolerant to mismatches in the central seed region that hybridizes with target RNAs^11–14^. However, whether the CRISPR-Cas13 system can discriminate circRNAs from their linear cognates with gRNAs targeting sequences spanning the BSJ, and if yes, the robustness and broad application of this approach on circRNA levels have remained unknown.

## RfxCas13d/BSJ-gRNA discriminates circRNAs from mRNAs

We first sought to identify the best effector for circRNA knockdown by screening Cas13 family proteins^11–15^. We synthesized the mammalian codon-optimized DNA sequence for each protein (LwaCas13a, PspCas13b, PguCas13b, RanCas13b, EsCas13d, AdmCas13d or RfxCas13d) and cloned them individually into an expression vector with a sequence for C-terminal monomeric superfolder GFP (msfGFP) to enhance protein stability (Extended Data Fig. 1c). Each Cas13 was expressed in 293FT cells (Extended Data Fig. 1d-e) and various knockdown efficiencies of *KARS* mRNA were observed when Cas13 and matched gRNAs were co-transfected into cells (Extended Data Fig. 1f), showing that these Cas13 proteins are capable of targeted RNA cleavage using specific gRNAs.

Next, we examined the knockdown efficiency and specificity of each Cas13 on two circRNAs, *circPOLR2A* (336 nt) and *circRTN4* (2,457 nt). We designed three gRNAs spanning BSJ (BSJ-gRNAs) for each circRNA (Fig. 1a). Most Cas13s (except LwaCas13a) could attenuate expression of both circRNAs but had no detectable effect on their linear cognate mRNAs (Fig. 1b; Extended Data Fig. 1g), suggesting that the Cas13 system could discriminate circRNAs from paired linear mRNAs by BSJ-gRNAs. Among all examined Cas13 constructs, RfxCas13d exhibited the highest knockdown efficiency (>80%) for both circRNAs, while linear *POLR2A* and *RTN4* mRNAs remained largely unchanged (Fig. 1b; Extended Data Fig. 1g). Northern Blot analysis further confirmed the dramatic knockdown of *circPOLR2A* by RfxCas13d/BSJ-gRNAs (Extended Data Fig. 1h). Importantly, a non-targeting (NT) gRNA did not direct Cas13 to circRNAs for cleavage (Fig. 1b; Extended Data Fig. 1g); overexpression of RfxCas13d or individual BSJ-gRNAs alone did not affect circRNA expression (Extended Data Fig. 1i-j). To rule out off-target effects of BSJ-gRNAs, we designed additional control gRNAs having half-sequences replaced by scrambled sequences (paired control) (Fig. 1c) or sequences from adjacent linear exons (gRNA-L) (Extended Data Fig. 1k) of *circPOLR2A* and *circRTN4*, respectively. None of these control gRNAs could direct RfxCas13d to degrade corresponding circRNAs (Fig. 1c and Extended Data Fig. 1k). As expected, however, control gRNAs (gRNA-L) targeting linear RNA exon-exon junctions led to decreased linear mRNA expression by RfxCas13d (Extended Data Fig. 1k).

**Fig. 1.**
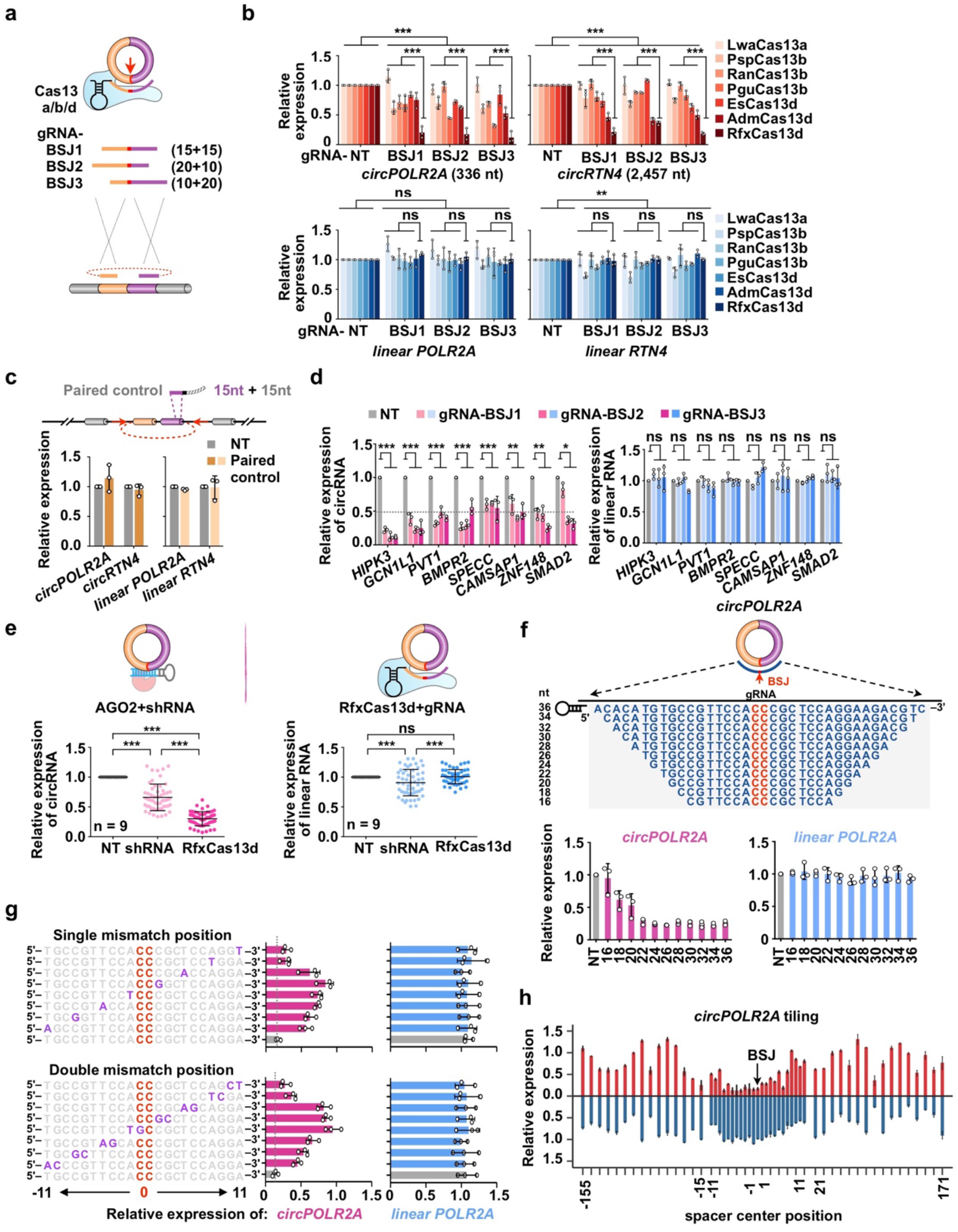
RfxCas13d/BSJ-gRNA discriminates circRNAs from mRNAs. **a.** Schematic of circRNA knockdown by Cas13 orthologues. Three guide RNAs targeting the back-splicing junction site (red arrow) were designed for each circRNA (BSJ-gRNAs). **b.** Evaluation of different Cas13-protein-mediated knockdown on circRNAs. Expression levels of two circRNAs, *circPOLR2A* and *circRTN4*, and their cognate mRNAs were detected by qRT-PCR in 293FT cells. NT, non-targeting guide RNA. **c.** A paired control gRNA with partial sequences replaced by scrambled sequences for *circPOLR2A* or *circRTN4* did not guide RfxCas13d to affect expression of circRNAs or mRNAs. **d.** The RfxCas13d/BSJ-gRNA system mediates specific and robust knockdown of randomly selected circRNAs, but not their cognate linear mRNAs in 293FT cells. **e.** The RfxCas13d/BSJ-gRNA system exhibits higher specificity and efficiency than shRNAs on circRNA knockdown. Knockdown of nine circRNAs with RfxCas13d/BSJ-gRNAs or position-matched shRNAs was compared. Each circRNA was targeted by two BSJ-gRNAs and two position-matched shRNAs; the expression of each cognate linear mRNA was also examined. **f.** Minimal length requirement of BSJ-gRNA spacer for efficient RfxCas13d knockdown. Top, lengths and sequences of gRNA spacers flanking the *circPOLR2A* BSJ site; bases flanking the BSJ site are shown in red. Bottom, knockdown efficiencies of *circPOLR2A* and linear *POLR2A* by each BSJ-gRNA and RfxCas13d were detected by qRT-PCR. **g.** Mismatch tolerance of RfxCas13d for circRNA targeting. Left, guides containing single or double mismatches at varying positions across spacer sequences for *circPOLR2A* are shown; mismatch positions are shown in purple; bases flanking the BSJ site are shown in red. Right, knockdown efficiencies of *circPOLR2A* and linear *POLR2A* by each BSJ-gRNA and RfxCas13d were detected by qRT-PCR. **h.** Efficiency and specificity of RfxCas13d for circRNA and cognate linear mRNA knockdown. Plots of knockdown efficiencies of 54 guides for *circPOLR2A*, including 23 guides tiled across the BSJ and 31 guides tiled on the overlap region of *circPOLR2A* and linear *POLR2A*. All transcript levels were normalized to *ACTB*, and values represent mean +/− SD with *n*=3, *: *P* < 0.05; **: *P* < 0.01; ***: *P* < 0.001; ns, not significant.

To explore the general applicability of RfxCas13d/BSJ-gRNAs for circRNA interference, we examined the knockdown efficiencies of another eight abundant circRNAs in human cells (Fig. 1d). Similar to *circPOLR2A* and *circRTN4*, expression of all examined circRNAs, but not paired linear mRNAs, decreased more than 50% when RfxCas13d and BSJ-gRNAs vectors were co-transfected into 293FT cells (Fig. 1d). In HeLa cells, we found that RfxCas13d was the most efficient Cas13 to specifically disrupt expression of circRNAs (Extended Data Fig. 2a-c).

siRNAs^5,16,17^ and shRNAs^3,4^ were used in loss-of-function (LOF) studies of circRNAs. We compared the specificity and knockdown efficiency of the RfxCas13d system and shRNAs in a position-matched manner for nine pairs of circular and linear cognate RNAs (Fig. 1e, top). Compared to AGO2/shRNA-mediated knockdown, RfxCas13d/BSJ-gRNA showed much higher efficiency of circRNA knockdown (RfxCas13d: mean 70%; shRNA: mean 34%) and much lower off-target rates on linear mRNAs for all examined pairs (Fig. 1e, bottom). Collectively, these results reveal that RfxCas13d is an effective tool for circRNA knockdown with high efficiency, specificity and generality.

## Characteristics of RfxCas13d/BSJ-gRNA-mediated circRNA knockdown

We characterized key features required for RfxCas13d/BSJ-gRNA-mediated circRNA knockdown. We first determined the minimal spacer length requirement for an efficient RfxCas13d targeting for circRNAs. We generated a series of BSJ-gRNAs with spacers ranging from 16~36 nt, symmetrically spanning the BSJ of *circPOLR2A* (Fig. 1f, top). Consistent with the previous *in vitro* cleavage assay of EsCas13d^11^, a minimal spacer of 18 nt was required for RfxCas13d to target *circPOLR2A*, but the highest knockdown efficiency was observed with 22 nt or longer spacers (Fig. 1f, bottom left). In all examined conditions, we observed that the expression of linear *POLR2A* mRNA was barely affected (Fig. 1f, bottom right), suggesting that recognizing the unique BSJ in *circPOLR2A* (considered as a mismatch ribonucleotide for the paired linear mRNA) in the spacer could effectively constrain RfxCas13d targeting to only the circRNA.

This notion was further confirmed by characterizing the mismatch tolerance of RfxCas13d for circRNA targeting. We introduced either single or double mismatches into BSJ-gRNAs and observed that any single- or double-mismatch strikingly impaired the knockdown efficiency of *circPOLR2A* by RfxCas13d (Fig. 1g). A central seed region of the BSJ-gRNA spacer was identified as being more sensitive to mismatches, extending from the position of −8 to 8 nt in a 22 nt long BSJ-gRNA spacer (Fig. 1g). As expected, expression of linear *POLR2A* mRNA remained unaltered under all conditions (Fig. 1g).

We evaluated the knockdown efficiency and specificity of RfxCas13d for circRNA and its paired linear mRNA by designing gRNAs tiled in 10-nt increments across the BSJ of *circPOLR2A* or *circPVT1* (Fig. 1h; Extended Data Fig. 3a-b). Only BSJ-gRNAs could guide RfxCas13d to knock down these circRNAs without affecting linear RNA, whereas other non-BSJ site containing gRNAs could direct RfxCas13d to degrade both circular and linear RNAs but to a lesser degree (Fig. 1h; Extended Data Fig. 3a-b).

We examined the position-effect of the BSJ-gRNA spacer for RfxCas13d-mediated circRNA knockdown at the single nucleotide resolution by tiling gRNAs across the BSJ of *circPOLR2A*. We observed that gRNA spacers with the BSJ in the center (−7 nt~7 nt spanning the BSJ site) exhibited high knockdown efficiency without affecting linear cognate RNAs (Fig. 1h; Extended Data Fig. 3c). Similar results were observed for two other circRNAs *circHIPK3* (1,099 nt) and *circRTN4* by tiling assays (Extended Data Fig. 3d-e). The advantage of the RfxCas13d system in mediating circular but not linear RNA interference (Fig. 1d, 1h; Extended Data Fig. 3) could be due to distinct biogenesis efficiencies, conformations and turnover rates between circular and linear RNAs^7,18^.

## RfxCas13d/BSJ-gRNA screens identify circRNAs that act in a cell-type specific manner in cell proliferation

Next, we set out to explore the feasibility of using the RfxCas13d/BSJ-gRNA system to perform large-scale, loss-of-function screens of circRNAs in human cells. We constructed a library of BSJ-gRNAs that target highly expressed circRNAs in human cells (Fig. 2a). Among top 10% highly expressed circRNAs with more than 3 copies in one line of HT29, HeLa, 293FT or H9 cells by using CIRCexplorer2^19^ (Extended Data Fig. 4a-b), 762 circRNAs were successfully included in the gRNA library (Fig. 2a; Supplementary Table 1). This pool of circRNAs contains the most highly expressed (>70%) circRNAs in examined human cell lines (Extended Data Fig. 4c) and has a representative length distribution compared to all expressed circRNAs (Extended Data Fig. 4d)

**Fig. 2.**
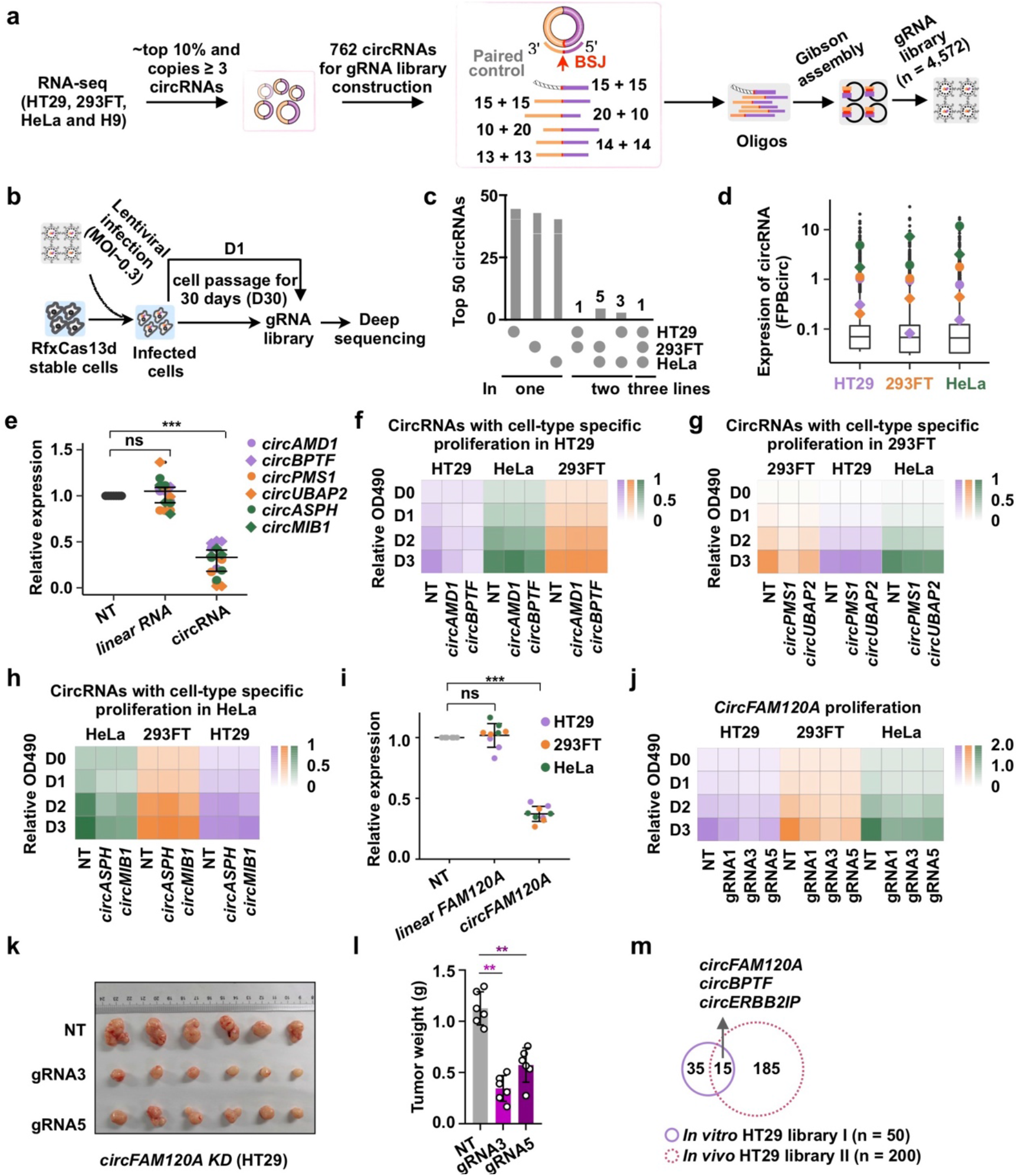
BSJ-gRNA library construction and circRNAs act in a cell-type specific manner in cell proliferation. **a.** Schematic of gRNA library design and construction. One paired control gRNA (n=762) and five BSJ-gRNAs (circRNA gRNAs, n=3,810) were designed for each candidate circRNA. In total, 4,572 gRNAs were included in the library. **b.** Screen of circRNAs necessary for cell growth and proliferation. The gRNA lentiviral library was individually delivered into HT29, 293FT and HeLa cells stably expressing RfxCas13d. Infected cells were collected at day 1 (D1) and day 30 (D30) after infection and genomic DNA was extracted for gRNA amplification and deep sequencing. **c.** Overlapping analysis of the top50 circRNA hits that potentially have impacts on cell growth by screens in HT29, 293FT and HeLa cells. **d.** Expression (shown by FPBcirc) of circRNAs that exhibit cell type-specific effect on cell proliferation in HT29, 293FT and HeLa cells. **e.** Knockdown of circRNAs shown in (d) by the RfxCas13d/BSJ-gRNA system in HT29, 293FT and HeLa cells, individually. Expression levels of circRNAs and linear mRNAs were detected by qRT-PCR. All transcript levels were relative to *ACTB*, and values represent mean +/− SD with *n*=3, ***: *P* < 0.001; ns, not significant. **f.** Cell type-specific growth effect of *circAMD1* and *circBPTF* in HT 29 cells, shown by MTT cell proliferation assays. **g.** Cell type-specific growth effect of *circPMS1* and *circUBAP2* in 293FT cells, shown by MTT cell proliferation assays. **h.** Cell type-specific growth effect of *circASPH* and *circMIB1* in HeLa cells, shown by MTT cell proliferation assays. **i.** Specific knockdown of *circFAM120A* by RfxCas13d/ BSJ-gRNAs in HT29, 293FT and HeLa cells. Expression levels of *circFAM120A* and linear *FAM120A* were detected by qRT-PCR. All transcript levels were relative to *ACTB*, and values represent mean +/− SD with *n*=3, ***: *P* < 0.001; ns, not significant. **j.** Knockdown of *circFAM120A* by RfxCas13d/ BSJ-gRNAs inhibits HT29, 293FT and HeLa cell growth and proliferation, as revealed by MTT cell proliferation assays. **k-l.** Knockdown of *circFAM120A* by RfxCas13d inhibits tumors growth. The image of tumors from nude mice injected subcutaneously with NT and *circFAM120A*-KD HT29 cells is shown in **(k)**, and the statistics of tumor weight is shown in **(l)**. **m.** Overlap of circRNA candidates between in cells (top 50, n=762) and *in vivo* (top 200, n=2,908) screens in HT29 cells. *CircFAM120A* was identified in both screens.

To construct the library, we designed five BSJ-gRNAs that all target sequences spanning the BSJ in each circRNA and with different lengths (26~30 nt) (Fig. 2a; Supplementary Table 1). Such BSJ-gRNAs were capable of mediating efficient and specific knockdown for *circPOLR2A*, *circHIPK3*, *circRTN4* and other examined circRNAs (Fig. 1; Extended Data Fig. 1-3). One paired control gRNA (Fig. 1c) was designed for each circRNA, which is more rigorous than scramble controls commonly used in screening studies (Fig. 2a). The gRNA library was constructed from synthetic oligo pools by Gibson assembly (Fig. 2a). Deep sequencing of enriched gRNAs amplified from the library showed a high representation and uniformity of gRNAs (Extended Data Fig. 4e).

We conducted screens to identify circRNAs that are required for cell proliferation. HT29, HeLa and 293FT cells stably expressing RfxCas13d were infected with the gRNA lentiviral library at a multiplicity of infection (MOI) of 0.3. These cells were cultured for 30 days followed by gRNA enrichment and deep sequencing (Fig. 2b). Read distributions of two biologically independent experiments showed a high correlation in HT29, HeLa and 293FT cells (Extended Data Fig. 4f-h). After 30 days of culture, some gRNAs were obviously depleted compared to paired control gRNAs in each cell line (Extended Data Fig. 4i-k), indicating these circRNAs might be important for cell growth. We ranked these circRNA candidates according to robust rank aggregation (RRA) scores (Extended Data Fig. 4l-n; Supplementary Table 2), calculated by MAGeCK that evaluates the statistical significance of individual gRNA abundance changes using a negative binomial model, and compared the ranks of gRNAs targeting each circRNA with a null model of uniform distribution^20^ (Extended Data Fig. 5a). A small RRA score indicates a strong negative selection of circRNAs during the screening. In general, we observed mild effects of circRNAs on cell proliferation. By considering low RRA score and high expression level, the top 50 circRNA hits in each cell line were obtained, which were potentially important for cell growth (Extended Data Fig. 4l-n; Supplementary Table 3).

We selected ~10 circRNAs in each cell line for validation by RfxCas13d/BSJ-gRNA. We began with one BSJ-gRNA of each circRNA for validation and the great majority of circRNAs could be successfully knocked down with expected cell growth inhibition in these individual types of cells (Extended Data Fig. 5b; Supplementary Table 4). Among them, *circHIPK3* and *circKLHL8* were previously reported to be necessary for cell proliferation by shRNA knockdown^21,3^. To exclude potential off-targets on their parental genes, three BSJ-gRNAs targeting *circHIPK3* or *circKLHL8* were designed to deplete these circles and all exhibited high knockdown efficacy and retarded cell proliferation in HeLa or 293FT cells (Extended Data Fig. 6a-c). These phenotypes were unlikely to have resulted from off-target effects on their parental genes which remained unchanged (Extended Data Fig. 6a-c). Their suppression of cell proliferation could be also reproduced by shRNA-mediated knockdown (Extended Data Fig. 6d-f), even though two shRNAs targeting *circKLHL8* displayed distinct effects (Extended Data Fig. 6d, 6f), indicating possible off-target effects of these shRNAs.

The majority of circRNA hits (>80%) were unique to only one examined cell type (Fig. 2c; Extended Data Fig. 7a). This cell-type specific mode of circRNA effects is unlikely due to different expression as they were expressed at similar levels in all three cell lines (Fig. 2d; Extended Data Fig. 7b). Further validation of circRNA hits by RfxCas13d/BSJ-gRNA (Fig. 2e) displayed the consistent observation that knockdown of these circRNAs exhibited a cell-type-specific inhibition of cell growth (Fig. 2f-h). Collectively, these results suggested the feasibility of using RfxCas13d/BSJ-gRNA for functional circRNA screening in human cells.

## *CircFAM120A* affects the proliferation of different cell types *in vitro* and *in vivo*

Our screens identified a previously unappreciated circRNA, *circFAM120A*, which is involved in proliferation of all examined cell types. *CircFAM120A* is highly expressed with over 20 copies per cell (Extended Data Fig. 8a). Knockdown of *circFAM120A* by RfxCas13d with three different BSJ-gRNAs all displayed a retarded cell proliferation in HT29, HeLa and 293FT cells (Fig. 2i-j; Extended Data Fig. 8b). The effect of *circFAM120A* on cell growth was also seen by shRNA-mediated knockdown (Extended Data Fig. 8c-f).

As RfxCas13d/BSJ-gRNAs could persistently knock down circRNAs, we examined the effect of *circFAM120A* on cell proliferation in nude mice by introducing HT29 cells stably transfected with the control or RfxCas13d-gRNA plasmids for *circFAM120A* knockdown. Consistently, two examined gRNAs targeting *circFAM120A* each showed that *circFAM120A* is important for cell growth *in vivo* (Fig. 2k-l). These results also suggest that applicability of RfxCas13d-gRNA for LOF circRNAs in xenograft models.

Next, to identify potential circRNA candidates involved in cell proliferation *in vivo*, we constructed a new library of BSJ-gRNAs that target 2,908 highly expressed circRNAs in nine human cells (Extended Data Fig. 9a-b; Supplementary Table 5). This pool of circRNAs contains the most highly expressed (>80%) circRNAs in examined human tissues (Extended Data Fig. 9c), thus representing a comprehensive pool for screening circRNAs *in vivo*. We carried out *in vivo* screens by transferring the BSJ-gRNA library-infected RfxCas13d-HT29 cells into nude mice and obtained xenografts 22 days after injection (Extended Data Fig. 9d). RNA-seq analyses revealed that the correlation between two biological repeats *in vivo* was lower than that in cell screening (Extended Data Fig. 9e). Among the top 200 circRNA hits identified from *in vivo* screens selected by RRA scores (Fig. 2m; Extended Data Fig. 9f), 15 hits appeared in the top 50 hits in HT29 cells screened by the small gRNA library (Fig. 2a, 2m; Extended Data Fig. 4l). *CircFAM120A* was among these overlap candidates.

## *CircFAM120A* promotes cell growth by up-regulating its parental gene translation

To gain insights into the mechanism of *circFAM120A* in cell proliferation, we performed transcriptomic analyses after *circFAM120A* knockdown by BSJ-gRNA or shRNA (Extended Data Fig. 10a). Compared to shRNA-mediated knockdown, the BSJ-gRNA-mediated knockdown showed a much higher correlation with gene expression (Extended Data Fig. 10b-c), further confirming the specificity and robustness of the RfxCas13d/BSJ-gRNA approach for knocking down circRNAs.

Gene ontology analysis of differentially expressed genes (DEGs) resulting from BSJ-gRNA-mediated *circFAM120A* knockdown revealed an enrichment of altered genes involved in cell proliferation and apoptosis (Extended Data Fig. 10d-f). For example, the expression of *EIF4EBP1*, *NUPR1* and *PCK2* with reported oncogenic function involved in the AKT pathway ^22–25^ was down-regulated upon *circFAM120A* knockdown (Extended Data Fig. 10g), supporting its role in promoting cell proliferation.

*CircFAM120A* was mainly localized in the cytoplasm (Extended Data Fig. 11a) with few enrichment on polyribosomes (Extended Data Fig. 11b), excluding the possibility of *circFAM120A* translation. As a few cytoplasmic circRNAs, such as *CDR1as* and *circHIPK3*, were found to act as miRNA sponges ^21,26^, we asked whether *circFAM120A* could act in a similar manner. Although predicted miRNA sites of Argonaut 2 (AGO2) binding could be identified computationally from published eCLIP datasets^26^ (Extended Data Fig. 11c-d), RNA Immuno-precipitation (RIP) assays with anti-AGO2 followed by examination of associated RNAs revealed no interaction between *circFAM120A* and AGO2; whereas other known positive controls including *CDR1as* and *circHIPK3* showed strong association (Extended Data Fig. 11e).

Given that *FAM120A* is an oncogenic gene related to the AKT pathway ^27^ and that AKT pathway was over-represented upon the loss of *circFAM120A* (Extended Data Fig. 10g). We next asked whether *circFAM120A* could promote cell proliferation by regulating the expression of its linear cognate *FAM120A* mRNA.

First, knockdown of *FAM120A* mRNA by shRNAs showed an inhibitory effect on 293FT and HT29 cell proliferation (Fig. 3b; Extended Data Fig. 12a-d). Second, knockdown of *circFAM120A* by RfxCas13d/BSJ-gRNA led to reduced FAM120A protein expression (Fig. 3c; Extended Data Fig. 12e) without affecting *FAM120A* mRNA level (Fig. 2i), indicating that this reduction might result from inhibited translation of *FAM120A* mRNA. Consistent with this possibility, polysome profiling showed a shift of linear *FAM120A* mRNA from polyribosomes to monoribosomes after *circFAM120A* knockdown by BSJ-gRNA (Fig. 3d-e; Extended Data Fig. 12f), indicating impaired translation. Third, transfection of FAM120A-overexpression vectors in *circFAM120A* deficient cells fully rescued the impaired cell proliferation (Fig. 3f). These results suggested that *circFAM120A* promotes cell growth by preventing translation of its cognate linear mRNA.

**Fig. 3.**
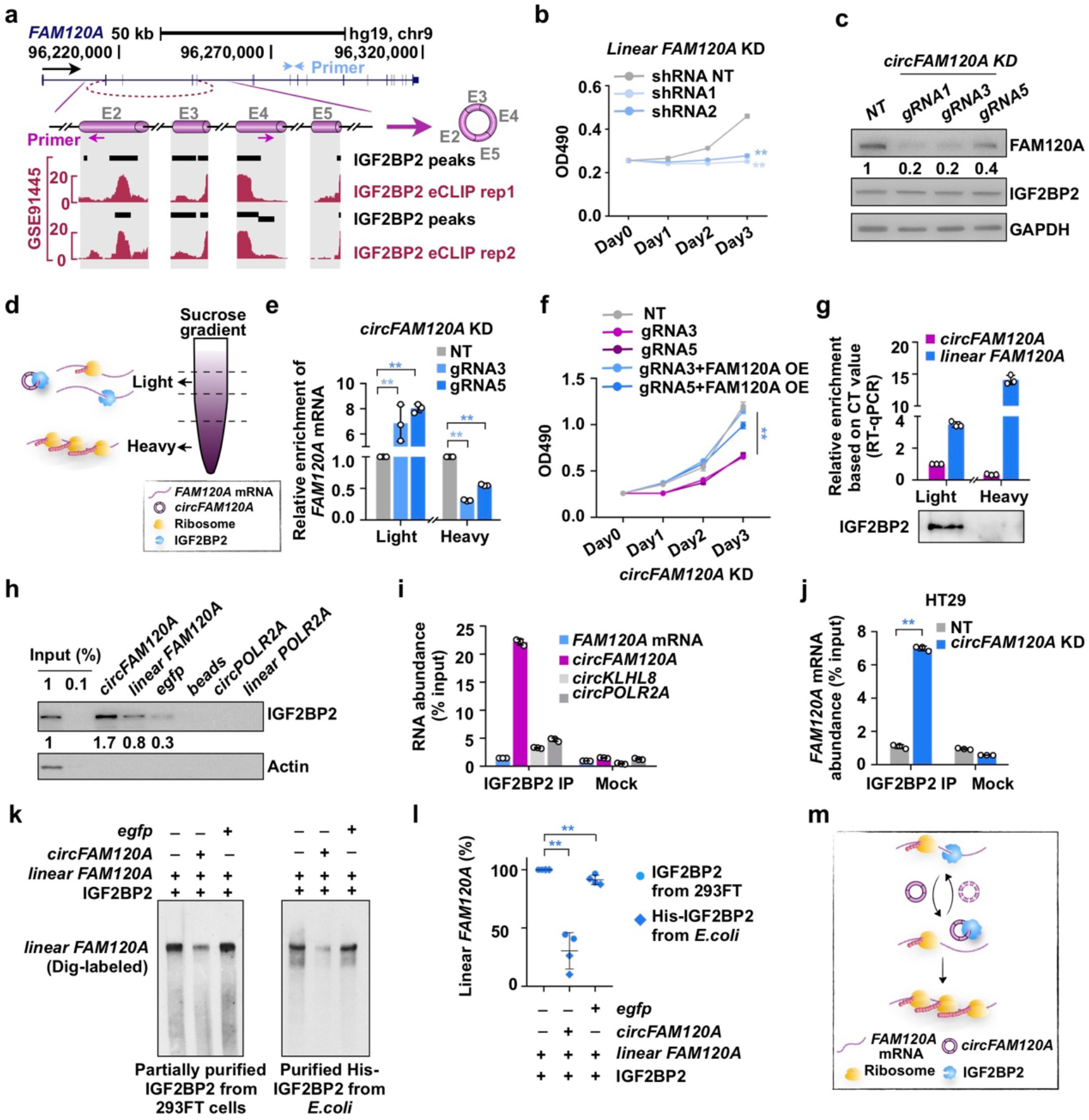
*CircFAM120A* promotes cell proliferation by preventing *FAM120A* mRNA from interacting with IGF2BP2 for efficient translation. **a.** Prediction of IGF2BP2-binding peaks in *circFAM120A*. Top, genomics locus and diagram of linear *FAM120A* and *circFAM120A* (shown as cylinders in magenta). Blue and magenta arrows indicate location of primer for linear *FAM120A* or *circFAM120A*, respectively. Bottom, IGF2BP2-binding peaks and wiggle-tracks of eCLIP-seq data in K562 cells (GEO, GSE91445) revealed that IGF2BP2 proteins were targeted on *circFAM120A* producing circularized exons. **b.** Knockdown of *FAM120A* mRNA by shRNAs inhibits 293FT cells growth and proliferation, as revealed by MTT cell proliferation assays. **c.** Knockdown of *circFAM120A* by RfxCas13d/ BSJ-gRNAs leads to reduced expression of FAM120A protein in 293FT cells. **d.** Schematic of the sucrose gradient used to segregate light and heavy fraction in polysome profiling assay. **e.** Knockdown of *circFAM120A* leads to reduced enrichment of linear *FAM120A* on polyribosomes (heavy fraction). Cytoplasmic extracts from NT or *circFAM120A*-KD 293FT cells, were loaded on 10%-45% sucrose gradients. Enrichments of *FAM120A* mRNA in light or heavy fraction was measured by qRT-PCR. All values represent mean +/− SD with *n*=3, **: *P* < 0.01. **f.** Overexpression of linear *FAM120A* mRNA rescues the impaired cell proliferation in *circFAM120A* deficient cells, as revealed by MTT cell proliferation assays. All values represent mean +/− SD with *n*=3, **: *P* < 0.01. **g.** The distribution of *circFAM120A*, linear *FAM120A* and IGF2BP2 protein in the light or heavy fraction of 293FT cells, as revealed by polysome profiling assays. **h.** *circFAM120A* prefers to bind to IGF2BP2 compared to linear *FAM120A* mRNA or other negative controls, as revealed by biotin-labeled RNA pull-down assays. **i.** IGF2BP2 perfers to bind *circFAM120A*, compared to linear *FAM120A* mRNA or other circRNAs, as revealed by IGF2BP2 RNA immunoprecipitation (RIP) in 293FT cells. The abundance of RIP-enriched RNAs was measured by qRT-PCR. All values represent mean +/− SD with *n*=3, **: *P* < 0.01. **j.** Knockdown of *circFAM120A* leads to increased interaction between cognate *FAM120A* mRNA and IGF2BP2 in HT29 cells. The abundance of *FAM120A* mRNA was measured by qRT-PCR, and values represent mean +/− SD with *n*=3, *: *P* < 0.05; **: *P* < 0.01. **k.** Addition of *circFAM120A* impaired IGF2BP2 and *FAM120A* mRNA interaction, as revealed by *in vitro* competition assay. The IGF2BP2 protein is immunoprecipitated from 293FT cells (left) or purified from the *E. coli* (right). **l.** Quantification of *in vitro* competition assays shown in (K). Data are shown as median and IQR. n.s., p > 0.05, **p < 0.01, Student’s t test. **m.** A proposed model of *circFAM120A* binding IGF2BP2 to prevent linear *FAM120A* mRNA from interaction with IGF2BP2 for an efficient translation to promote cell growth. See text for details.

## *CircFAM120A* prevents *FAM120A* mRNA from binding to IGF2BP2, leading to efficient translation

To explore how does *circFAM120A* regulate *FAM120A* mRNA translation, we screened RBPs associated with the *circFAM120A*-producing locus from published eCLIP databases (https://www.encodeproject.org/), and identified IGF2BP2, an RNA binding protein with reported function in cell proliferation by suppressing mRNA translation ^28,29^ (Fig. 3a). Polysome profiling revealed distinct distributions of IGF2BP2, *circFAM120A* and the cognate linear *FAM120A* mRNA (Fig. 3g; Extended Data Fig. 12g). In agreement with its inhibitory function in translation, >95% IGF2BP2 was localized to the ribosome-free fraction, where both circular and linear *FAM120A* RNAs were absent. The majority of *circFAM120A* presented in the light monoribosome fraction, where <5% IGF2BP2 was enriched (Extended Data Fig. 12g); in contrast, only ~10% of *FAM120A* mRNAs were localized to monoribosomes and most *FAM120A* mRNAs were associated with polyribosomes in the heavy ribosome fraction (Fig. 3g; Extended Data Fig. 12g). These findings indicated that *circFAM120A* might compete with its cognate mRNA for IGF2BP2 binding in monoribosomes and that the IGF2BP2-unbound-*FAM120A* mRNA could be engaged with polyribosomes for translation. To confirm this hypothesis, we performed a series of RNA pull-down and IGF2BP2 RIP assays. The strong and specific association of IGF2BP2 with *circFAM120A*, but not other circRNAs, was observed (Fig. 3h-i). Knockdown of *circFAM120A* led to enhanced interaction between linear *FAM120A* mRNA and IGF2BP2 in HT29 cells (Fig. 3j) and 293FT cells (Extended Data Fig. 12h). *In vitro* competition assays with IGF2BP2 immunoprecipitated from 293FT cells or purified from *E. coli* both showed that the addition of *circFAM120A* strikingly impaired IGF2BP2 and *FAM120A* mRNA interaction (Fig. 3k-l). Thus, although sharing the same sequences with *circFAM120A*, the linear mRNA showed much weaker interaction with IGF2BP2 than *circFAM20A* (Fig. 3h-i). These results are consistent with a role of *circFAM120A* in promoting human cell proliferation by preventing *FAM120A* mRNA from binding to IGF2BP2 for efficient translation (Fig. 3m).

## Application of the RfxCas13d/BSJ-gRNA system to identify key circRNAs important for the development of mouse embryos

It has been shown that circRNAs are abundant in early mouse embryos from the zygote to blastocyst stages^30^. However, it remains unknown whether any circRNA is important during development. To identify such circRNAs, we first explored whether the RfxCas13d/BSJ-gRNA system is applicable to mouse embryos. To do this, we microinjected purified RfxCas13d mRNAs and guide RNAs targeting *Kras* and *Brg1* mRNAs into zygotes, followed by the examination of knockdown efficiency of each mRNA and defects in the preimplantation stage embryos (Fig. 4a). RfxCas13d/gRNAs displayed efficient knockdown of both *Kras* and *Brg1* mRNAs (Extended Data Fig. 13a). Only knockdown of *Brg1*, but not *Kras*, led to reduced blastocyst formation as previously reported^31,32^ (Extended Data Fig. 13b-d), suggesting the specificity and non-toxicity of using this system to study RNA function in mouse embryos.

**Fig. 4.**
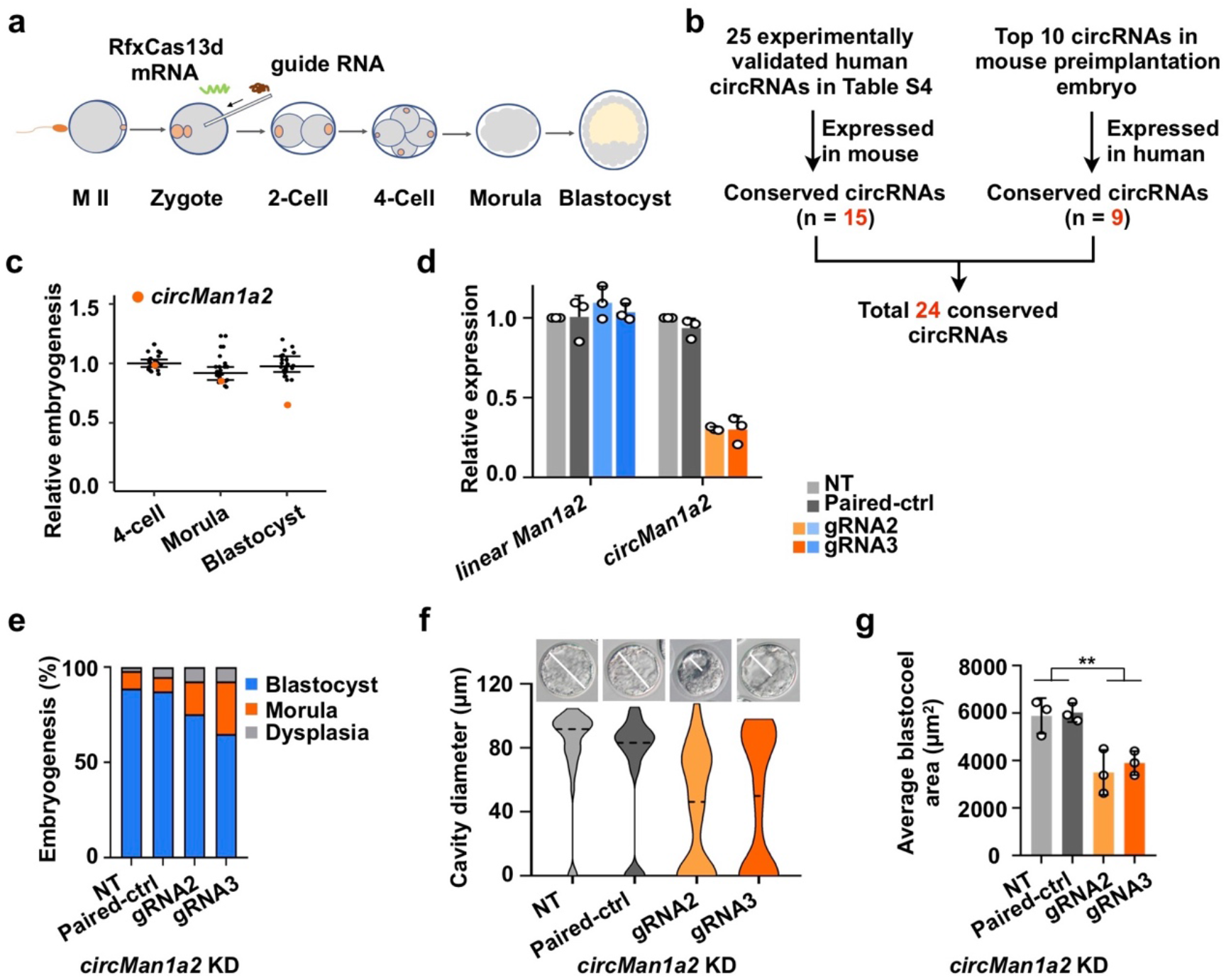
Screening circRNAs with functional potential during mouse preimplantation development with RfxCas13d/BSJ-gRNA. **a.** Schematic of RfxCas13d mRNA (green curve) and gRNA (brown curve) microinjection into mouse zygotes for RNA knockdown during preimplantation development. **b.** A schematic workflow to show the strategy of selecting 24 circRNAs tested in the preimplantation development of mouse embryos. The expression data of circRNAs in mouse preimplantation embryo from GEO: GSE53386^30^. **c.** The effect of embryogenesis of 24 circRNA candidates knockdown individually by RfxCas13d/BSJ-gRNA during mouse preimplantation development. **d.** Knockdown of *circMan1a2* by RfxCas13d/BSJ-gRNAs in zygotes. Expression levels of *circMan1a2* and linear *Man1a2* were detected by qRT-PCR. All transcript levels were relative to *Gapdh*, and values represent mean +/− SD with *n*=3. **e.** Knockdown of *circMan1a2* in zygotes led to reduced blastocyst formation 72h after microinjection of RfxCas13d mRNA and the corresponding BSJ-gRNAs into mouse zygotes. **f-g.** Cavity diameter **(f)** and area **(g)** of blastocysts were measured at 72h after microinjection of RfxCas13d mRNA and *circMan1a2* BSJ-gRNAs into mouse zygotes. Bright-field images showed the representative cavity diameter of blastocyst in each group **(f)**. The blastocoel area was calculated as length × width. All values represent mean +/− SD with *n*=3, **: *P* < 0.01.

Next, we applied the RfxCas13d/BSJ-gRNA system to screen potentially functional circRNAs during preimplantation development of mouse embryos. We selected a pool of 24 mouse circRNAs containing 15 circRNAs obtained from our screens in human cell lines (Fig. 2c) with mouse homolog circRNA expression in examined mouse cells and another nine highly expressed and conserved circRNAs from mouse preimplantation embryos^30^ (Fig. 4b). We confirmed their high expression during mouse preimplantation development (Supplementary Table 6). Initial screening by injecting RfxCas13d mRNA with one gRNA each targeting 24 individual circRNAs into more than 30~50 mouse zygotes identified one circRNA (*circMan1a2*) as a potential candidate important for mouse embryo preimplantation development (Supplementary Table 6).

*CircMan1a2* is a conserved circRNA that is derived from exons 2-6 of its parental gene in mouse or in human (Extended Data Fig. 14a). To exclude any potential off-target, one paired control and another BSJ-gRNA for *circMan1a2* were also tested. Consistent with results obtained in human cell lines (Fig. 1 and Extended Data Fig. 1-2), RfxCas13d efficiently targeted *circMan1a2* with BSJ-gRNAs with little effect on its cognate mRNA in mouse embryos (Fig. 4d). Importantly, microinjection of RfxCas13d mRNA and BSJ-gRNAs targeting *circMan1a2* into mouse zygotes led to retarded embryonic development as shown by decreased blastocyst formation rate (Fig. 4e) and smaller blastocoel according to the diameter and area of cavity (Fig. 4f-g; Extended Data Fig. 14b). As controls, knockdown other circRNAs, such as *circDcbld2*, had no obvious phenotype (Extended Data Fig. 14c; Supplementary Table 7). Together, these screens indicated potentially regulatory roles of circRNAs in the preimplantation stage of mouse embryos.

## Discussion

Due to the complete sequence overlap between back-spliced RNA circles and their linearly spliced RNA isoforms except BSJs, the functional study of individual circRNAs has been impeded by the lack of tools uniquely targeting these circles. We have shown that the RfxCas13d/BSJ-gRNA system achieves precise and robust circRNA-specific knockdown without disturbing their linear cognate mRNAs (Fig. 1a-d) and that this system is applicable at both individual and genome-wide levels (Fig. 2a-b). The Cas13 approach is unique to study LOF of circRNAs, which are derived from the middle exons of genes^33^. There are at least two reasons that account for this. First, the overall conformation of circRNAs is more stable and rigid than that of linear RNAs^18^. Thus, once targeted by gRNA and Cas13 in a circRNA, such rigid structures would be continuously to be targeted by this system; whereas in a linear RNA, the highly dynamic and flexible folding status theoretically make it difficult for effective targeting. Second, back-splicing efficiency is extremely slow, which is less than 1% of that of the canonical splicing in cells^7^. Once cleaved by Cas13, the newly produced circRNAs cannot compensate for the loss of circRNAs, leading to a consistent reduction of circRNAs by this system.

Screening of circRNAs involved in cell growth in different human cells suggests the cell-type specific manner of action of circRNAs, identifying only a few that likely act in examined human cells (Fig. 2a-h). This observation is consistent with a recent finding that knockdown of long noncoding RNAs can affect gene expression in a cell type-specific manner as revealed by CRISPRi-based genome-scale screening^34^. In the case of circRNAs, considering their species-^35^, tissue- and cell type-specific expression patterns ^36,7,37^, our results indicate that the great majority of circRNAs may not play generic housekeeping roles but some may act in a regulatory fashion in a cell type-specific manner.

Although most act in the cell-specific manner, it is worthwhile noting that our screens identified one commonly-expressed oncogenic circRNA, *circFAM120A*, which is derived from the *FAM120A* gene with reported roles in cancer^27^. *CircFAM120A* promotes cell proliferation both *in vitro* and *in vivo* (Fig. 2i-l). *CircFAM120A* is largely localized to monoribosomes (Fig. 3d, 3g), where it interacts with translation inhibitor IGF2BP2^28,29^ to prevent the binding between IGF2BP2 and *FAM120A* mRNA, facilitating the IGF2BP2-unboud mRNA translation (Fig. 3g-m).

The RfxCas13d/BSJ-gRNA system could also be used to effectively and robustly disturb linear and circular RNA expression in mouse embryos (Fig. 4a-d; Supplementary Tables 6-7). Screening a small pool of conserved circRNAs in mouse embryos identified *circMan1a2* that is important for mouse embryo preimplantation, suggesting previously unknown functional relevance of circRNAs in the early animal development. Generation of circRNA loss-of-function mouse models without affecting their parental mRNA expression will be a necessity to prove the physiological importance of endogenous circRNAs. Nevertheless, future application of the RfxCas13d/BSJ-gRNA system will likely enable the discovery of circRNAs that participate in other biological processes.

## Acknowledgments

We thank Chen and Yang laboratories for discussion. This work was supported by the Chinese Academy of Sciences (CAS) (XDB19020104), the National Natural Science Foundation of China (NSFC) (31725009, 31821004, 31730111, 31730062), the Shanghai Municipal Commission for Science and Technology (19411951800) and the Howard Hughes Medical Institute International Program (55008728).

## Author contributions

L.-L.C. supervised and conceived the project. L.-L.C., L.Y., J.L., S.-Q.L., X.L., L.Z., and W.X. designed experiments. L.Z. performed circRNA screening in mouse embryos supervised by J.L; S.-Q.L., X.L., L.Z., S.-M.C., L.-Z.Y., J.-L.Z., X.G., S.-K.G. and J.W. performed all other experiments; W.X., Y.-N.L., L.Y. preformed computational analyses. L.-L.C. and L.Y. wrote the paper with input from S.-Q.L., X.L., W.X and J.L.

## Competing interests

The authors declare that they have no competing financial interests.

## Supplemental Information

### Materials and Methods

#### Cell culture

HT29, HeLa and 293FT cells were purchased from the American Type Culture Collection (ATCC; http://www.atcc.org) or ThermoFisher, respectively, and were originally authenticated using STR profiling. HT29 cells were maintained in RPMI 1640, HeLa (human female origin) and 293FT cells (human fetus origin) were maintained in DMEM, and all supplemented with 10% Fetal Bovine Serum (FBS) and 0.1% penicillin/streptomycin. We maintained cell lines at 37°C in a 5% CO_2_ cell culture incubator and tested all cell lines routinely to exclude mycoplasma contamination.

#### Plasmid construction and transfection

For construction of Cas13 expression vectors, human codon-optimized DNA sequences of seven Cas13 orthologues (LwaCas13a, PspCas13b, PguCas13b, RanCas13b, EsCas13d, AdmCas13d and RfxCas13d) were synthesized in Genscript company and individually cloned into the p23-phage vector containing sequences for a C-terminal fused msfGFP and a Flag tag. DNA sequence of RfxCas13d was further cloned into the p23-phage vector containing sequences for a C-terminal fused mCherry and a Flag tag (p23-RfxCas13d-mcherry-flag). For construction of guide (g)RNA expression vectors, DNA sequences for gRNAs were synthesized and cloned into PUC19 or pLKO.1-TRC containing direct repeats of each Cas13 correspondingly. For construction of shRNA expression vectors, DNA sequences of shRNAs were synthesized and individually cloned into the pLKO.1-TRC vector. DNA sequences of all gRNAs and shRNAs are listed in Supplementary Table 6.

Plasmid transfection was carried out using Lipofectamine 3000 Reagent (ThermoFisher) according to manufacturer’s protocols. About 80%–90% transfection efficiency was achieved in HeLa and 293FT cells. Cells were harvested upon 48h after transfection for further RNA expression analysis.

#### Lentivirus production and cell infection

To produce lentiviral particles, 5 × 10^6^ 293FT cells in a 10-cm dish were co-transfected with 10 μg expression vector of interest, 7.5 μg psPAX2 vector and 3 μg pMD2.G vector. The supernatant containing viral particles was harvested upon 48h and 72h after transfection and passed through Millex-GP Filter Unit (0.22 mm pore size, Millipore). Viral particles were then concentrated 100-fold by sucrose gradient ultracentrifugation, resuspended in PBS containing 0.1% BSA, and stored at −80°C till use. To infect HeLa or 293FT cells with lentivirus, cells were incubated with culture medium containing 10 μL concentrated lentivirus and 5 mg/mL polybrene (Sigma). Infected cells were treated with puromycin for several days to increase knockdown efficiency.

#### Generation of stable cell lines with RfxCas13d expression

p23-RfxCas13d-mcherry-flag vector was infected into HeLa or 293FT cells by lentivirus for stable cell line generation. Single clones with red fluorescence were selected and validated by WB using anti-Flag antibodies.

#### RNA isolation and qRT-PCR

Total RNAs from cultured cells were extracted with Trizol (Life Technologies) according to the manufacturer’s protocol. RNAs were treated with DNase I (Ambion, DNA-freeTM kit). cDNAs were reverse transcribed with PrimeScript RT Reagent Kit (Takara) according to the manufacturer’s protocol for qPCR. *ACTB* was examined as an internal control for normalization. Primers for all qRT-PCR assays were listed in Supplementary Table 6.

#### Northern Blotting (NB)

NB was performed according to the manufacturer’s protocol (DIG Northern Starter Kit, Roche). Digoxigenin (Dig)-labeled antisense riboprobes were made using RiboMAX Large Scale RNA Production Systems (Promega). In brief, 5 μg total RNAs were resolved on denaturing urea polyacrylamide gel, transferred to nylon membrane (Roche) and UV-crosslinked using standard manufacturer’s protocols. Membrane was then hybridized with specific Dig-labeled riboprobes. Primers for NB probes to detect *circPOLR2A* were listed in Supplementary Table 6.

#### Western Blotting (WB)

Cells were harvested after treated and resuspended in lysis buffer (150mM NaCl, 1% NP-40, 0.5% sodium deoxycholate, 0.1% SDS, 50mM Tris, pH8.0, and protease inhibitors) for 10 min. After centrifugation, supernatants of cell lysates were resolved on 10% SDS-polyacrylamide gel and analyzed by WB with anti-Flag (Sigma), anti-FAM120A, anti-IGF2BP2, anti-IGF1BP1, anti-GAPDH or β-actin (Sigma) antibodies.

#### Cell proliferation assays

To validate the observed effect of a specific circRNA on cell proliferation, BSJ-gRNAs targeting BSJ site of the circRNA were individually infected into HeLa or 293FT cell lines stably expressing RfxCas13d. Cells with specific circRNA-knockdown were obtained by puromycin selection for several days. Knockdown efficiencies of circRNAs by RfxCas13d/BSJ-gRNAs or shRNAs were measured prior to performing cell proliferation assays.

Cell proliferation was detected by an MTT assay that measures OD_490_ according to CellTiter 96 AQueous One Solution Cell Proliferation Assay (Promega), or by Ensight® Multimode Plate Reader (PerkinElmer) that measures the occupied surface area of cells to indicate confluency. 5 × 10^3^ HeLa or 293FT cells were seeded into each well of 96-well dishes, then cultured at 37°C in a 5% CO_2_ cell culture incubator for indicated time points. Cell proliferation at Day 0 was detected after cells were seeded for 5h to allow their attachments to the plate. Live cells at D1, D2 and D3 were detected by MTT or by Ensight® Multimode Plate Reader. Data were normalized to the value of Day 0.

#### Subcutaneous xenograft tumor Models

All animal procedures were performed under the ethical guidelines of the Shanghai Institute of Biochemistry and Cell Biology, Chinese Academy of Sciences, CAS Center for Excellence in Molecular Cell Science. For subcutaneous xenograft tumor formation assay, 4 to 5-week-old male BALB/c-nu/nu mice (five mice per group) were injected subcutaneously with 1 × 10^6^ cells. When mice were sacrificed, the tumors were collected and weighed.

#### Subcellular fractionation

Nuclear and cytoplasmic fractionation was carried out as described ^38^ with slight modifications. Briefly, 3 × 10^6^ cells were washed with PBS and collected by centrifugation at 1,000 rpm for 2 min at room temperature. Cells were resuspended in 300 μL lysis buffer (DPBS containing 1% NP-40, 0.5% sodium deoxycholate, 5 mM EDTA, 1mM DTT, 1mM PMSF, 2mM RVC, 15% glycerol, 1 x proteinase inhibitor cocktail), incubated on ice for 5 min and then examined under an optical microscope to ensure that membranes of most of cells were disrupted. Cells were centrifuged at 4,000 x g for 1min at 4°C and the supernatant was saved as cytoplasmic fractionations. The precipitated nuclei were further washed once with lysis buffer, resuspended in 300 μL lysis buffer, sonicated and centrifuged at 13,000 rpm for 10 min at 4°C. The supernatant was saved as nuclear extracts.

#### *In vitro* RNA transcription, circularization and purification

In vitro RNA transcription assay was carried out as previous described ^18^ with slightly modification. 1μg PCR-amplified T7-DNA fragments were incubated with 2 μL T7 RNA polymerase enzyme and 0.5 mM dNTPs. 2 mM GMP was supplemented in the reaction to produce 5’-monophosphate RNA for *FAM120A* circularization. The reaction was carried out for 2 hr at 37°C, followed by DNase I treatment for 30 min at 37°C to remove DNA templates. Transcribed RNAs were precipitated with ethanol, washed with 70% ethanol and resuspended in RNase-free water.

For *FAM120A* circularization, 50 μg transcribed *FAM120A* were incubated with T4 RNA ligase 1 (NEB) in 500 μL reaction for overnight at 16°C according to the manufacturer’s protocol. Circularized *FAM120A* were treated with RNase R as described ^33^. Circularized and linear *FAM120A* were precipitated with ethanol and resolved on denaturing urea polyacrylamide gel. Then visualized by Ethidium bromide staining. The corresponding to circular RNA band was excised and circular RNAs were purified.

#### Circular/linear RNA pull-down

Biotinylated linear and circular RNAs were heated for 5 min at 65°C in PA buffer (10 mM Tris HCl pH 7.5, 10 mM MgCl_2_, 100 mM NH_4_Cl) and slowly cooled down to room temperature. Then, 5 × 10^6^ cells were resuspended with 1 mL binding buffer (10 mM HEPES pH 7.0, 50 mM KCl, 10% glycerol, 1 mM EDTA, 1 mM DTT, 0.5% Triton X-100, heparin 0.3 mg/mL), sonicated and centrifuged at 13,000 rpm for 10 min at 4°C. The supernatant was pre-cleared with Streptavidin Dynabeats (Invitrogen) for 30 min at room temperature, followed by incubation with folded RNAs for 30 min and with beads for 10 min. Retrieved proteins were subjected to WB.

#### Circular/ linear RNAs competition assay

Synthesized linear and circular *FAM120A* were heated for 5 min at 65°C in RNA-folding buffer (10mM HEPES and 10 mM MgCl_2_) and slowly cooled down to room temperature. 0.5 μg folded linear *FAM120A* and 0.5 μg *circFAM120A* were added to incubate with 1 μg His-tagged IGF2BP2 for 2 hr at 4°C in 0.2 mL binding buffer (50 mM HEPES at pH 7.0, 150 mM NaCl, 10 mM MgCl_2_, 0.1 mM DTT, 0.5 mM PMSF, 2 mM RVC). *circFAM120A* or linear *FAM120A* binding with His-tagged IGF2BP2 were collected by anti-His antibody and extracted with Trizol (Life technologies) according to the manufacture’s protocol. Northern blot used to test the relative binding abundance of *circFAM120A* or linear *FAM120A*.

#### RNA Immunoprecipitation (RIP)

Cells growing in 15 cm dishes were rinsed twice with ice-cold PBS, harvested in 10 mL ice-cold PBS and then centrifuged at 1,000 rpm for 5 min at 4°C. Cell were resuspended in 1 mL RIP buffer (50 mM Tris, pH 7.4, 150 mM NaCl, 0.5% Igepal, 1mM PMSF, 1 × protease inhibitor cocktail (Roche) and 2 mM VRC) and subjected to three rounds of gentle sonication. Cell lysates were centrifuged at 12,000 rpm for 15 min at 4°C and the supernatants were precleared with 15 mL Dynabeads Protein G (Invitrogen) to get rid of non-specific binding. Then, the precleared lysates were used for IP with anti-AGO2 or anti-IGF2BP2 antibodies. IP was carried out for 2 hr at 4°C. The beads were washed three times with high salt buffer and two times with the RIP buffer, followed by extraction with elution buffer (100mM Tris, pH 6.8, 4% SDS, and 10mM EDTA) at room temperature for 10 min. One-third of the eluted sample was used for WB and the remaining was used for RNA extraction. The RNA enrichment was assessed by qRT-PCR. Primers are listed in Supplementary Table 6.

#### Ribosome fractionation

*circFAM120A* KD and control cells were exposed to cycloheximide (100 μg/mL) for 15 min, then 2 × 10^7^cells were lysed in 500 μL lysis buffer (20 mM Tris pH 7.4, 15 mM MgCl_2_, 200mM KCl, 1% Triton X-100, 100 μg/mL cycloheximide, 1mM DTT, 1 mg/mL heparin and 40 U/mL RNasin (Promege)), and centrifuged at 13,000 rpm at 4°C for 10 min. For fractionation, the lysates were loaded on 10%-45% sucrose gradients and sequenced by ultracentrifugation with a SW41 rotor (Beckman) at 36,000 rpm at 4°C for 2.5 hr. Linear sucrose gradients were prepared with a Gradient Master (Biocomp) as indicated by the manufacturer. The distribution of ribosomes on the gradients was recorded at 254 nm by BIOCOMP Piston Gradient Fractinator equipped with BIO-RAD ECONO UV Monitor.

#### Zygote injection and *in vitro* embryo culture

Around 8-week old B6D2F1 (C57BL/6 X DBA/2) female mice were superovulated and mated with the male B6D2F1 mice. 21-23 hr later, fertilized embryos were collected from oviducts. Afterwards, Rfx-Cas13d mRNAs (100 ng/μl) and gRNA (100 ng/μl) were mixed and injected into the cytoplasm of fertilized eggs in a droplet of HEPES-CZB medium containing 5 μg/ml cytochalasin B (CB) using a FemtoJet microinjector (Eppendorf) with constant flow settings. The injected embryos were cultured in KSOM with amino acids at 37°C under 5% CO2 in air until hatching blastocyst stage by 4 days.

#### Candidate circRNA selection and gRNA design for RfxCas13d/BSJ-gRNA library

For library of 762 circRNAs construction, we applied CIRCexplore2^19^ to identify circRNAs in multiple cell lines (HT29, 293FT, HeLa and H9 cells). In total, 11,796 circRNAs were identified. To obtain highly expressed circRNAs, we selected the top 10% highly expressed circRNAs (1,490) that present in at least one line of HT29, HeLa, 293FT or H9 cells. According to the reported 6 copies of *circPOLR2A* per HeLa cell ^18^, we calculated *circPOLR2A* and other circRNA copy numbers in each cell line by the FPBcirc value of each circRNA from RNA-seq. In total, 1,179 circRNAs were selected by expression at ≥ 3 copies in at least one cell line. Among them, 762 circRNAs (65%) were successfully designed with BSJ-gRNAs and included in the gRNA library (Supplementary Table 1).

For each 762 circRNA, we designed five BSJ-gRNAs that all target sequences spanning BSJ with different lengths (15nt + 15nt, 20nt + 10nt, 10nt + 20nt, 14nt + 14nt and 13nt + 13nt). In addition, one paired control gRNA (random 15nt + matched 15nt), in which their half-sequences were replaced by scrambled sequences, was designed for each circRNA and was included in the gRNA library (Supplementary Table 1).

For library of 2,908 circRNAs construction, we applied CIRCexplore2^19^ to identify circRNAs in multiple cell lines (HT29, 293FT, HeLa, H9, HepG2, K562, PA1, SH-SY5Y and FB cells). In total, 50,911 circRNAs were identified. To obtain highly expressed circRNAs, we selected the top 800 highly expressed circRNAs (3,177) of each cell lines. Among them, 2,908 circRNAs were successfully designed with BSJ-gRNAs and included in the gRNA library (Supplementary Table 2).

For each 2,908 circRNAs, we designed three BSJ-gRNAs that all target sequences spanning BSJ with different lengths (15nt + 15nt, 20nt + 10nt and 10nt + 20nt). In addition, one paired control gRNA (random 15nt + matched 15nt), in which their half-sequences were replaced by scrambled sequences, was designed for each circRNA and was included in the gRNA library (Supplementary Table 2).

#### Construction of a RfxCas13d/BSJ-gRNA library

We created a library targeting 762 circRNAs and 2,908 circRNAs with 4,572 gRNAs and 11,832 gRNAs as mentioned above. Oligonucleotides containing gRNA sequences were synthesized in Genscript company. Then, primers matching flanking sequences of oligonucleotides were used for the amplification to create 60-bp homologies with BsmBI digested gRNA-expressing backbone ^39^. The amplified DNA products were assembled into the BsmBI digested gRNA-expressing backbone using Gibson assembly method and were transformed into *E. coli* DH5α Electro-Cells (Takara) by electroporation to obtain the library plasmids. The lentivirus of the gRNA library was produced by co-transfection of library plasmids with psPAX2 and pMD2.G plasmids into 293FT cells using the Lipofectamine 3000 Reagent.

#### Screening for circRNAs with functional potential using the RfxCas13d/gRNA library

To screen circRNAs that are important for cell proliferation, RfxCas13d stable cells were infected with the gRNA lentiviral library at a multiplicity of infection (MOI) of 0.3. After selection by puromycin for one day, 2 × 10^6^ infected cells were harvested as D1 samples. And an aliquot of 2 × 10^6^ infected cells were cultured and passaged for 30 days as D30 samples. Two biologically independent experiments were performed both in HeLa and 293FT cells. All samples were used for DNA extraction, gRNA enrichment by PCR amplification and deep sequencing.

To screen circRNAs whose loss may confer resistance of cells to doxorubicin (DOX) treatment, RfxCas13d stable 293FT cells were infected with the gRNA lentiviral library at a MOI of 0.3. After selection by puromycin for one day, 2 × 10^6^ infected 293FT cells were cultured and passaged for 7 days. Then infected cells were treated with DMSO or DOX (1μM) for 2 days, survived cells were collected and used for DNA extraction, gRNA enrichment by PCR amplification and deep sequencing. Two biologically independent experiments were performed.

Deep sequencing of amplified PCR products containing gRNA sequences was conducted by Illumina NextSeq 500 at CAS-MPG Partner Institute for Computational Biology Omics Core, Shanghai, China.

#### Computational analysis of circRNA screens

Adapters at 3’ end and 5’ end in the raw paired-end sequencing datasets were removed by cutadapt (1.16) using the following adapter sequences:

R1: -a TTTTTTAAGCTTGGCGTAACTAGATCT -m 15,
-g CCCTACCAACTGGTCGGGGTTTGAAAC -m 15;
R2: -a GTTTCAAACCCCGACCAGTTGGTAGGG -m 15,
-g AGATCTAGTTACGCCAAGCTTAAAAAA -m 15.

Processed reads were aligned to gRNA library sequences with Bowtie (v1.1.2, parameters: -v 3 -m 1 -k 1). Only uniquely mapped reads of each sample were calculated.

The read number for each unique gRNA was normalized as previously reported^40^. Counts of gRNAs from two biological replicates were assessed by Pearson correlation coefficient (PCC). Average of normalized reads from two biological replicates were used as the normalized read count for each gRNA.

RRA score of each circRNA was calculated by MAGeCK that evaluates the statistical significance of individual gRNA abundance change using a negative binomial model ^20^. CircRNAs were ranked by RRA scores with a null model of a uniform distribution. A small RRA score indicates a strong selection of circRNAs during the screening. Fifty circRNAs with the lowest RRA scores were selected, and their normalized gRNA reads of paired-controls were unchanged between D1 and D30 samples. Finally, circRNAs with top 50 RRA score in HT29, 293FT or HeLa cells were selected as potential candidates. See Extended Data Fig. 5.

#### Polyadenylated RNA library preparation, deep sequencing and analysis

Polyadenylated [poly(A)+] RNA enrichment was performed as described^41^. Poly(A)+ RNA-seq libraries were prepared using Illumina TruSeq Stranded mRNA Sample Prep Kits and subjected to deep sequencing with Illumina NextSeq 500 at the CAS-MPG Partner Institute for Computational Biology Omics Core, Shanghai, China. RNA-seq sequencing read quality was evaluated by FastQC (v0.11.5).

Deep sequencing datasets were mapped with TopHat (TopHat v2.0.12, parameters: – microexon-search -g 1 -a 6 -m 2) and aligned to GRCh37/hg19 human reference genome with the UCSC Genes annotation (Human: hg19 knownGene.txt updated at 2013/06/30). Gene expression of linear mRNAs was determined by FPKM (Fragments Per Kilobase of transcript per Million mapped fragments). The maximum FPKM of expressed transcript was selected to represent the expression level of each given gene. Refseq genes with FPKM ≥ 1 at least in one sample were selected for comparison (Human hg19 refFlat.txt updated at 2017/04/09), and mean of two replicates was calculated as the gene expression level. Fold change (FC) between gRNA (or shRNA) KD and RfxCas13d NT (or shRNA NT) was used to determine upregulated (FC ≥ 1.5), unchanged (0.667 < FC < 1.5) and downregulated (FC ≤ 0.667) genes.

#### Gene Ontology (GO) of Differentially Expressed Genes (DEGs)

DEGs after *circFAM120A* knockdown by RfxCas13d gRNA were identified for GO. Ninety seven DEGs were manually clustered with https://www.genecards.org/ (GeneCards) and https://www.ncbi.nlm.nih.gov/pubmed/ (PubMed). Annotated GO terms (http://amigo.geneontology.org/amigo) were employed to classify *circFAM120A*-affected genes based on their functions with a manual check^42^.

#### Conservation of circRNAs analyses

Conservation of circRNAs analysis pipeline according to previous publications^35^ with some modification. In brief, sequences of back-spliced exons were extracted and used LiftOver tool (http://genome.ucsc.edu/cgi-bin/hgLiftOver, parameters: -bedPlus D 3 -tab -minMatch D 0.1 -minBlocks D 1) to identify orthologous coordinates between human (genome version: hg19) and mouse (genome version: mm10). In addition, an expressed circRNA identified in mouse orthologous locus without nucleotide difference and also expressed in mouse was suggested as a conserved circRNA. Similar strategy was also applied to mouse circRNAs.

#### Statistical analyses

*P* values < 0.05, < 0.01 and < 0.001 were marked by 1 asterisk, 2 asterisks or 3 asterisks, respectively. Statistically significant difference was assessed using Wilcoxon test with R platform (R v.3.5.1) and statistical significance was set at *P* < 0.05. To evaluate the relevant correlation between two datasets, Pearson correlation coefficient (PCC) was performed with R platform (R v.3.5.1). Statistically significant difference for cell proliferation was assessed using T test, and statistical significance was set at *P* < 0.05.

## Data availability

All sequencing datasets have been deposited in the NCBI GEO (GSEXXX) and National Omics Data Encyclopedia (OEPxxxxxx).

## Supplemental Figures and Figure Legends

**Extended Data Fig. 1.**
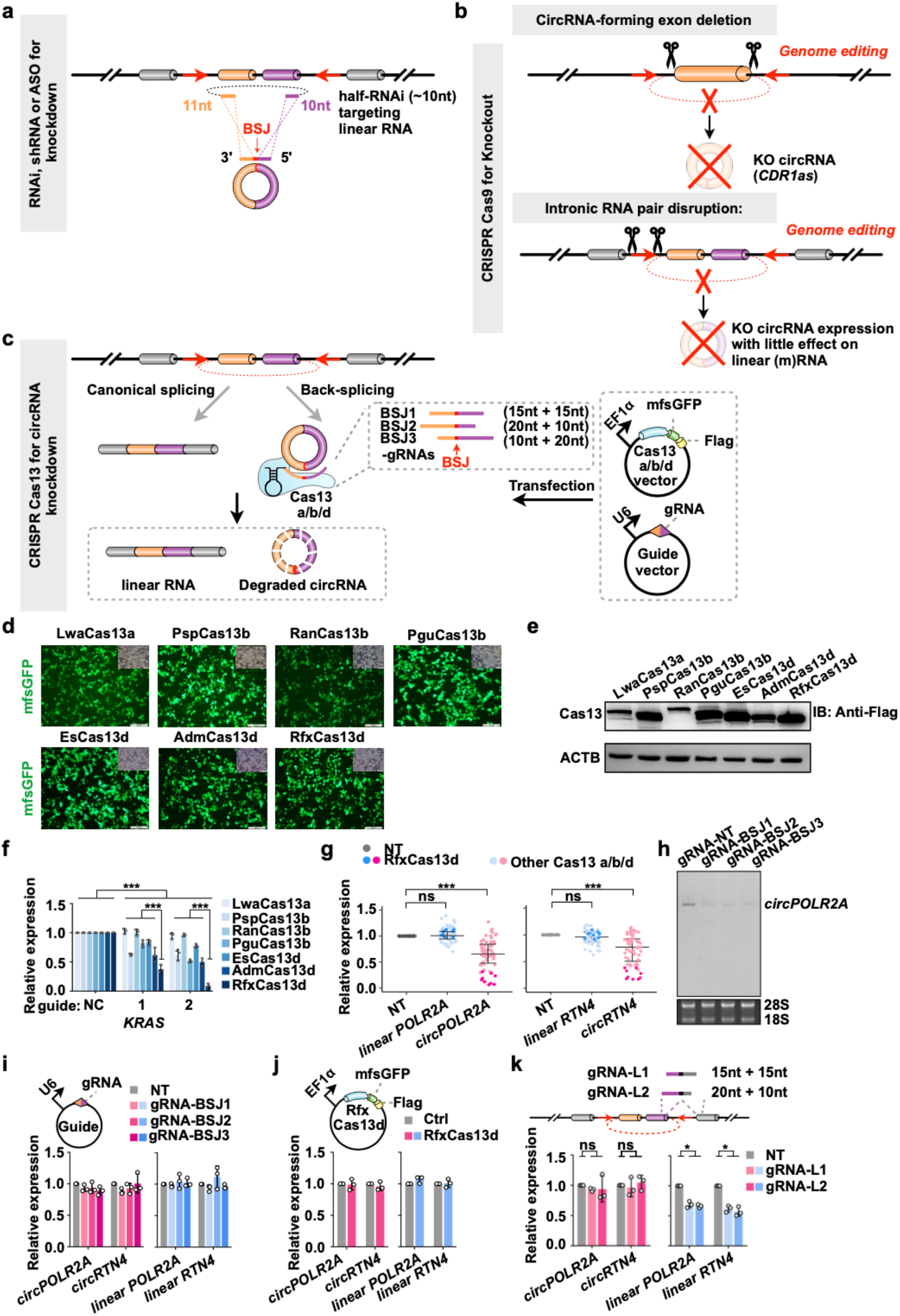
Currently applied interference technologies for circRNA expression and evaluation of RfxCas13d knockdown efficiency and specificity. **a**, Strategy of circRNA knockdown by RNAi machinery, such as RNAi, shRNA and ASO. The BSJ is a featured site for specific targeting of circRNA. Note, even half-RNAi (~10nt) can target the linear cognate RNA. **b**, Strategies of circRNA knockout by CRISPR/Cas9. Top, genome editing is used to directly remove the circRNA-formed exons; bottom, genome editing destroys the intronic RNA pair to block circRNA expression with little effect on the linear cognate RNA. **c**, Schematic of circRNA knockdown by Cas13 orthologues. Cas13a/b/d sequences were individually cloned into the vector with sequences for a C-terminal monomeric superfolder GFP (msfGFP) and a Flag-tag. Three guide RNAs targeting the back-splicing junction site (BSJ, red arrow) were designed for each circRNA (BSJ-gRNAs). Expression levels of circRNAs and their cognate linear mRNAs were detected by qRT-PCR. NT, non-targeting guide RNA. **d**,**e**, Expression levels of Cas13 proteins were detected by msfGFP fluorescence (**d**) and western blot (**e**) in 293FT cells after 48 h transfection. **f**, Comparison of different Cas13 proteins-mediated *KRAS* knockdown efficiencies with two position-matched guides revealed that RfxCas13d is the best effector^15^. **g**, Evaluation of different Cas13-protein-mediated knockdown on circRNAs. Expression levels of two circRNAs, *circPOLR2A* and *circRTN4*, and their cognate mRNAs were detected by qRT-PCR in 293FT cells. NT, non-targeting guide RNA. **h**, Northern blot confirmed the *circPOLR2A* knockdown by RfxCas13d/BSJ-gRNAs. **i**,**j**, Overexpression of individual BSJ gRNAs (**i**) or RfxCas13d (**j**) alone did not affect *circPOLR2A* and *circRTN4* expression. **k**, gRNAs with partial sequences replaced by adjacent linear exon led to linear but not circular RNA knockdown. Top, schematic design of gRNAs that target exons with partial sequences overlapped with circRNAs, and the remaining sequences overlapped with their linear mRNA exons (gRNA-L). Bottom, knockdown efficiencies of *circPOLR2A* and *circRTN4* as well as their corresponding linear RNAs under the treatment of each control gRNA-L and RfxCas13d were detected by qRT-PCR. All transcript levels were normalized to *ACTB*, and values represent mean +/− SD with n=3, *: *P* < 0.05; **: *P* < 0.01; ***: *P* < 0.001; ns, not significant.

**Extended Data Fig. 2.**
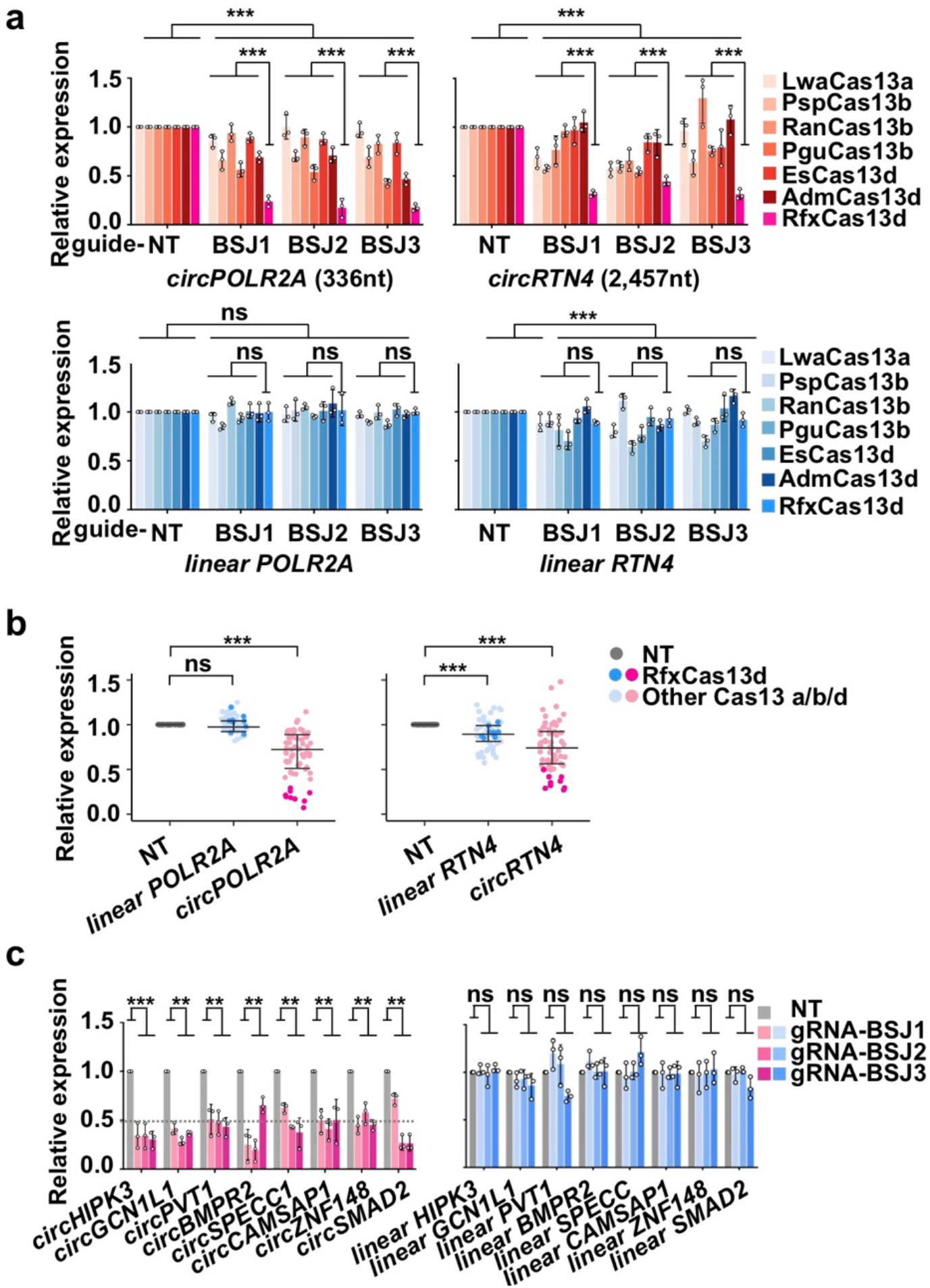
RfxCas13d is the best effector to mediate circRNA knockdown in HeLa cells. **a,b**, Evaluation of different Cas13 proteins mediated knockdown on circRNAs (*circPOLR2A* and *circRTN4*) and their cognate mRNAs in HeLa cells revealed that RfxCas13d is the best effector for the circRNA-specific knockdown. **c**, The RfxCas13d/BSJ-gRNA system mediates specific and robust knockdown of randomly selected circRNAs, but not their cognate mRNAs in HeLa cells. All transcript levels were relative to *ACTB*, and values represent mean +/− SD with n=3, **: *P* < 0.01; ***: *P* < 0.001; ns, not significant.

**Extended Data Fig. 3.**
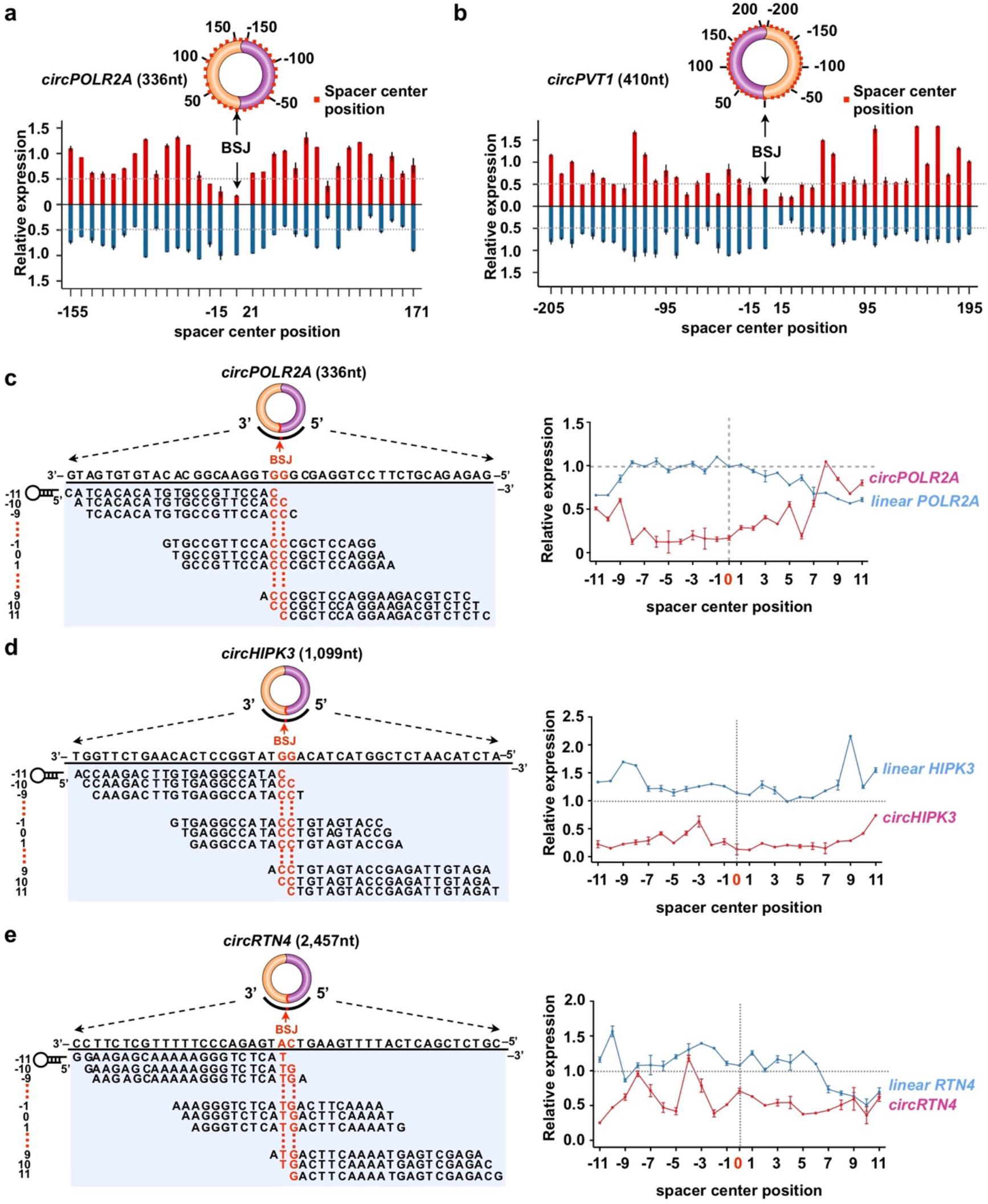
RfxCas13d arrayed screening of BSJ site of targeted circRNAs. **a,b**, Efficiency and specificity of RfxCas13d for circRNA and cognate linear mRNA knockdown. Top, schematic of gRNAs targeting *circPOLR2A* **(a)** and *circPVT1* **(b)** tiled 10-nt increments away from the BSJ site. Bottom, knockdown efficiencies of circRNA and linear RNA by each gRNA and RfxCas13d were detected by qRT-PCR. **c-e**, Arrayed knockdown screen of 23 guides evenly tilled across BSJ of *circPOLR2A* **(c)**, *circHIPK3* (**d**) and *circRTN4* (**e**). Position-effect of BSJ-gRNAs for RfxCas13d-mediated knockdown of *circPOLR2A* **(c)**, *circHIPK3* (**d**) and *circRTN4* (**e**) at the single nucleotide level. Twenty-three guides tiled across the BSJ of *circPOLR2A* **(c)**, *circHIPK3* (**d**) and *circRTN4* (**e**) are listed. Knockdown efficiencies of *circPOLR2A* **(c)**, *circHIPK3* (**d**) or *circRTN4* (**e**) and linear *POLR2A*, linear *HIPK3* or linear *RTN4* by each BSJ-gRNA were detected by qRT-PCR. All transcript levels are relative to *ACTB*, and values represent mean +/− SD with n=3.

**Extended Data Fig. 4.**
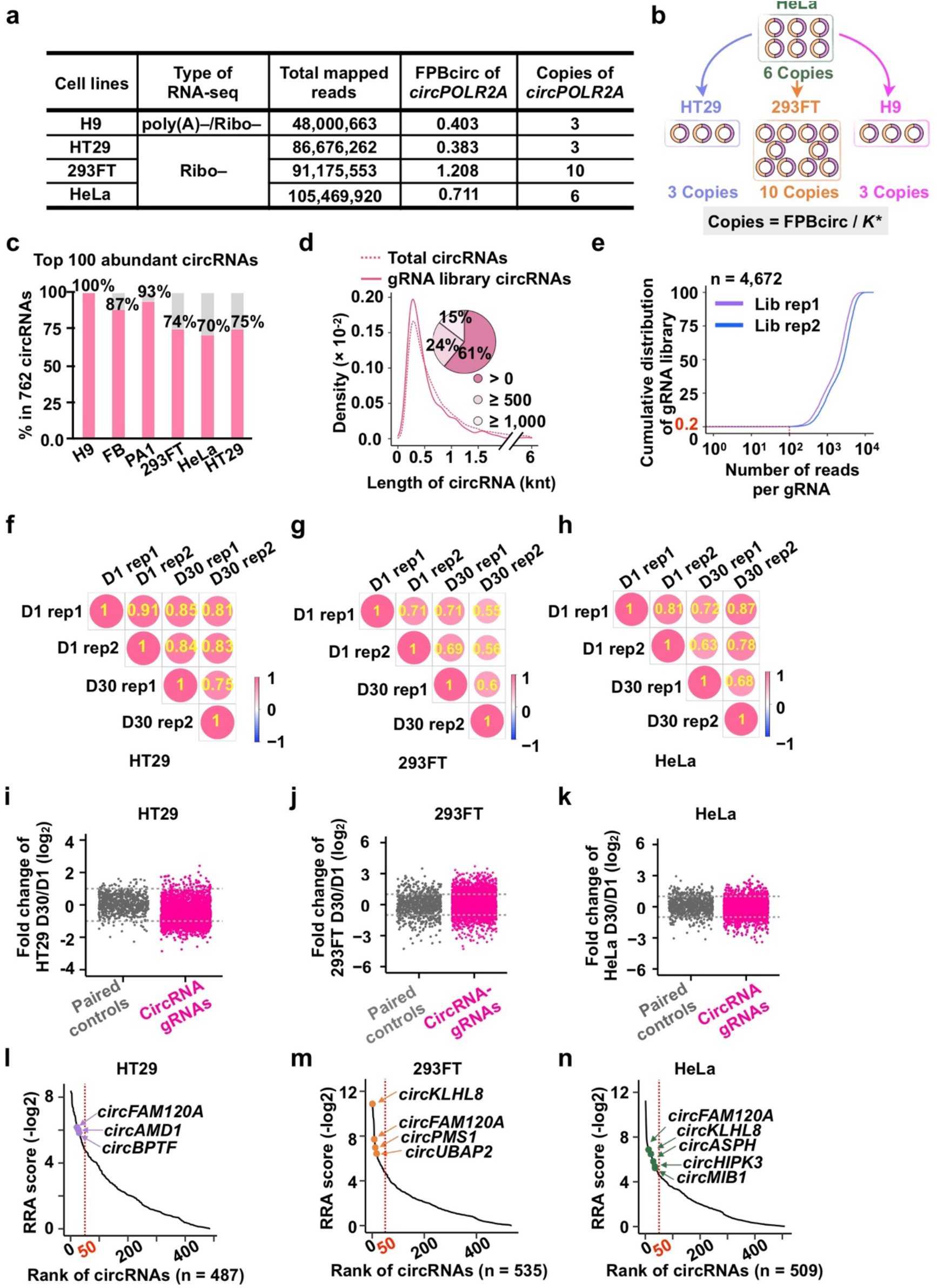
Characterization of the BSJ-gRNA library targeting 762 circRNAs and identification of negatively selected candidate circRNAs important for cell growth. **a**, Calculation of circRNA copy numbers (using *circPOLR2A* as an example) in H9, HT29, HeLa and 293FT cells. According to the FPM of *circPOLR2A* and other circRNAs in H9, HT29, HeLa and 293FT cells, and the known six copies of *circPOLR2A* per HeLa cell ^18^, we calculated the copy number of all circRNAs, respectively. For example, there are three copies of *circPOLR2A* per H9 or HT29 cells. **b**, Schematic of circRNA copy number calculation in H9, HT29, HeLa and 293FT cells. **c**, Representation of 762 candidate circRNAs designed with BSJ-gRNAs in the library in different cell lines. More than 70% of the top 100 abundant circRNAs in each indicated cell line were included in the 762 candidate circRNAs. **d**, Matched length distribution of 762 candidate circRNAs (solid line) and total circRNAs (dashed line). Density curve and pie chart show that more than 80% of 762 candidate circRNAs and total circRNAs are less than 1,000nt. **e**, Cumulative distribution of the number of reads per gRNA of constructed libraries. The red line indicates that less than 0.2% of gRNAs are covered by less than 100 reads. **f-h**, The Pearson correlation coefficient (PCC) between replicates (rep) of D1 and D30 samples in HT29 (**f**), 293FT (**g**) and HeLa cells **(h)**. Two biologically independent experiments were performed at D1 and D30 in each cell line. **i-k**, Scatter plot of fold change of paired controls and circRNA BSJ-gRNAs between D1 and D30 samples in HT29 **(i)**, 293FT **(j)** and HeLa cells **(k)**. The grey dashed lines indicate 2 or 0.5 fold change, respectively. **l-n**, Rank of negatively selected candidate circRNAs by RRA scores in HT29 **(l)**, 293FT **(m)** and HeLa cells **(n)**. Fifty circRNAs with the lowest RRA scores were sub-grouped by red dashed line.

**Extended Data Fig. 5.**
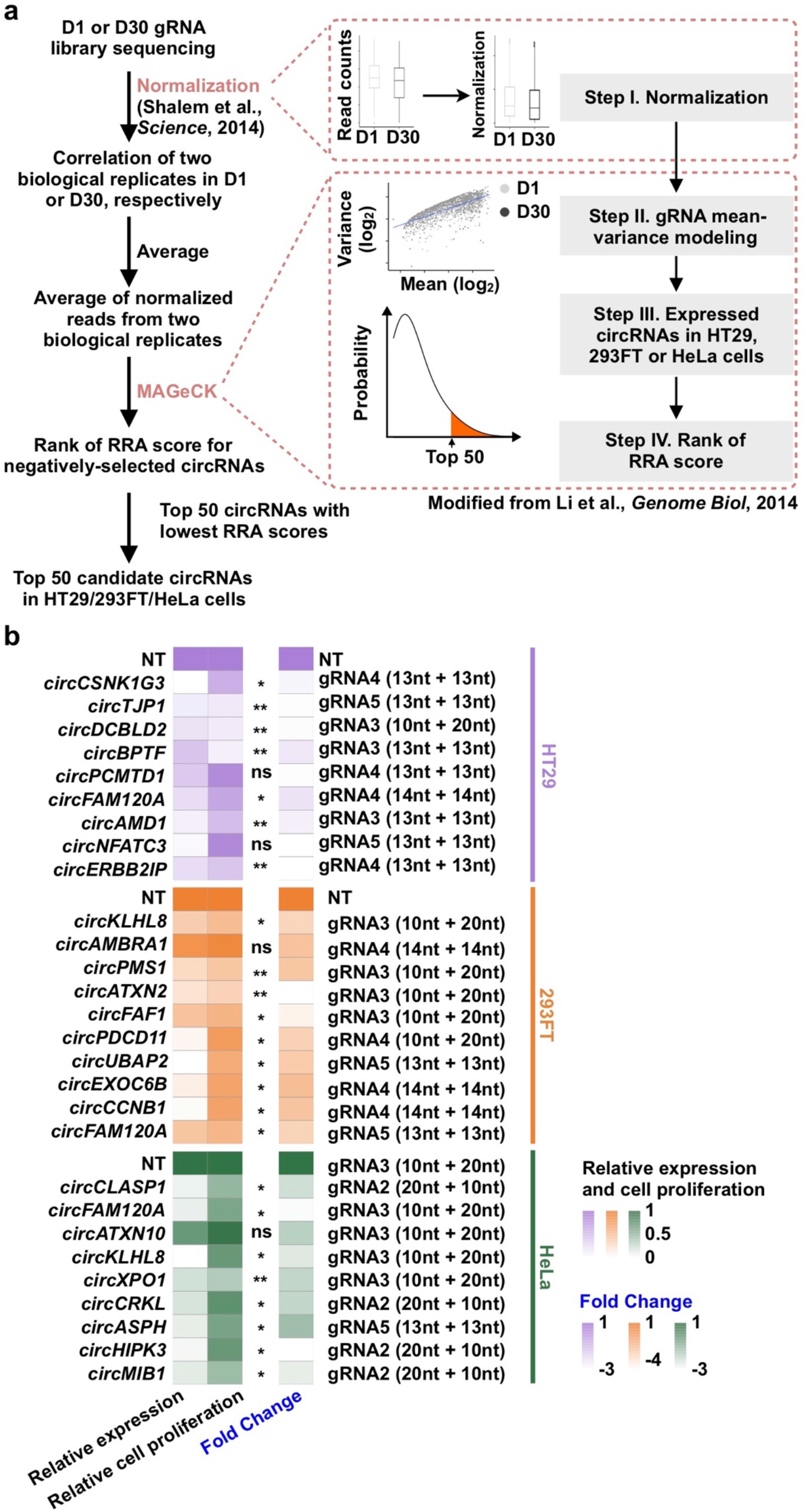
Validation of 25 candidate circRNAs by the RfxCas13d system in cell proliferation. **a**, Schematic of circRNA candidates identification. Raw read counts were normalized according to previous publication^40^. Since two biological replicates are well correlated, the averaged read counts were used for the subsequent analysis. RRA scores were calculated by the MAGeCK algorithm to determine top negatively selected circRNAs with low RRA scores. After that, expressed circRNAs in HT29, 293FT and HeLa cells were selected for future analysis. **b**, Heatmap display of the relative knockdown efficiency, cell proliferation and fold change of candidate circRNAs in HT29 (purple), 293FT (orange) and HeLa (green) cells with single BSJ-gRNA that targets each candidate circRNA.

**Extended Data Fig. 6.**
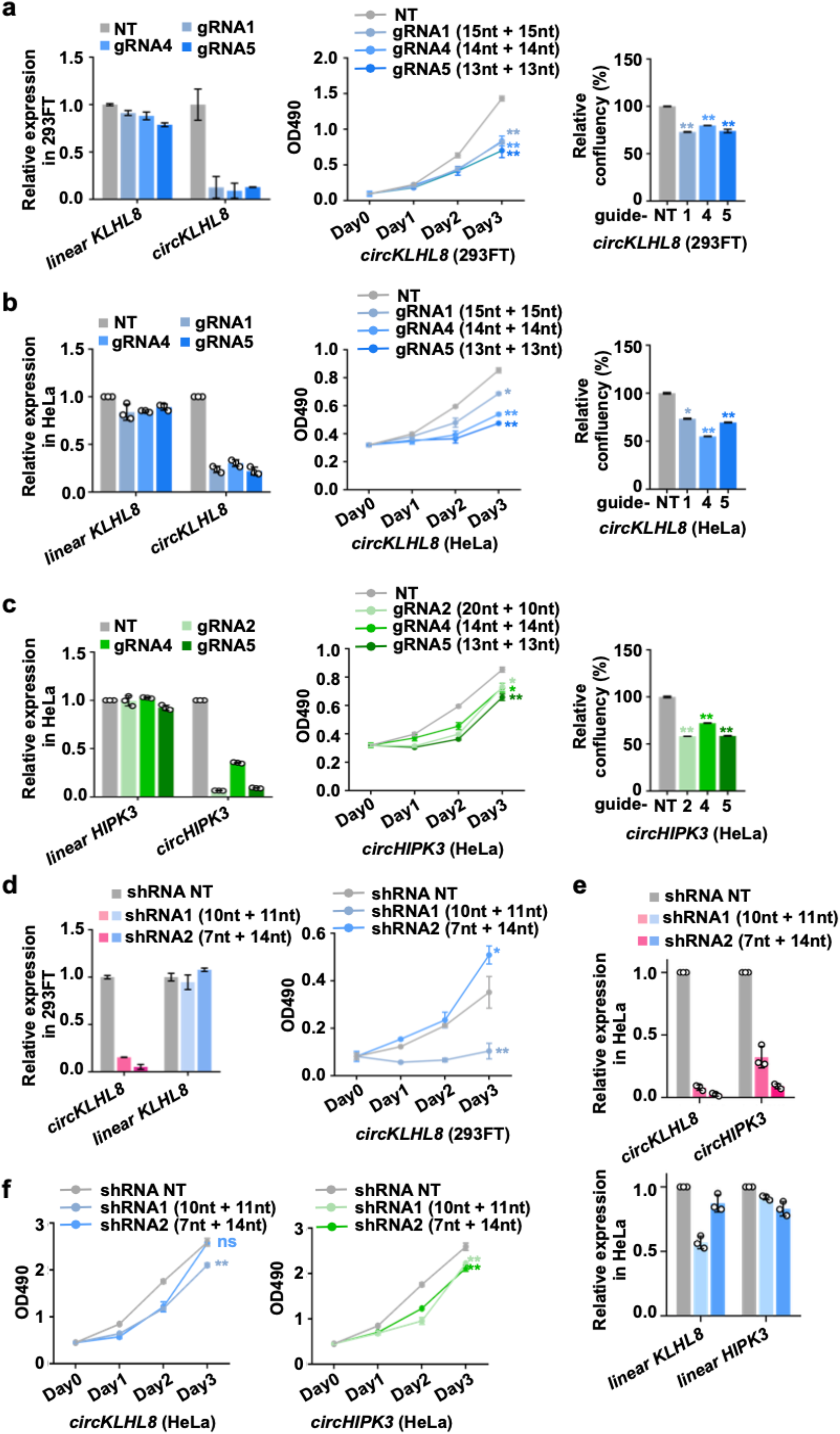
Validation of *circKLHL8* and *circHIPK3* by the RfxCas13d system and shRNAs in cell proliferation. **a,b**, Knockdown of *circKLHL8* by RfxCas13d inhibited cell proliferation in 293FT **(a)** and HeLa cells **(b)**, as revealed by MTT cell proliferation assay (Middle) and cell confluency calculated by the surface area occupied by cells (Right), the knockdown efficiency is showed in left. **c**, Knockdown of *circHIPK3* by RfxCas13d inhibited cell proliferation in HeLa cells, as revealed by MTT cell proliferation assay (Middle) and cell confluency calculated by the surface area occupied by cells (Right), the knockdown efficiency is showed in left. All transcript levels were normalized to *ACTB* by qRT-PCR, and values represent mean +/− SD with *n*=3, values represent mean +/− SD with *n*=3, *: *P* < 0.05; **: *P* < 0.01; ns, not significant. **d**, Knockdown of *circKLHL8* by shRNA inhibited cell proliferation in 293FT cells. The left is knockdown efficiency, and the right is cell proliferation. **e**, Knockdown of *circKLHL8* and *circHIPK3* by shRNA in HeLa cells. **f**, Individual knockdown of *circKLHL8* or *circHIPK3* by shRNA inhibited cell proliferation in HeLa cells, as revealed by MTT cell proliferation assay. All transcript levels were normalized to *ACTB* by qRT-PCR, and values represent mean +/− SD with *n*=3, *: *P* < 0.05; **: *P* < 0.01; ns, not significant.

**Extended Data Fig. 7.**
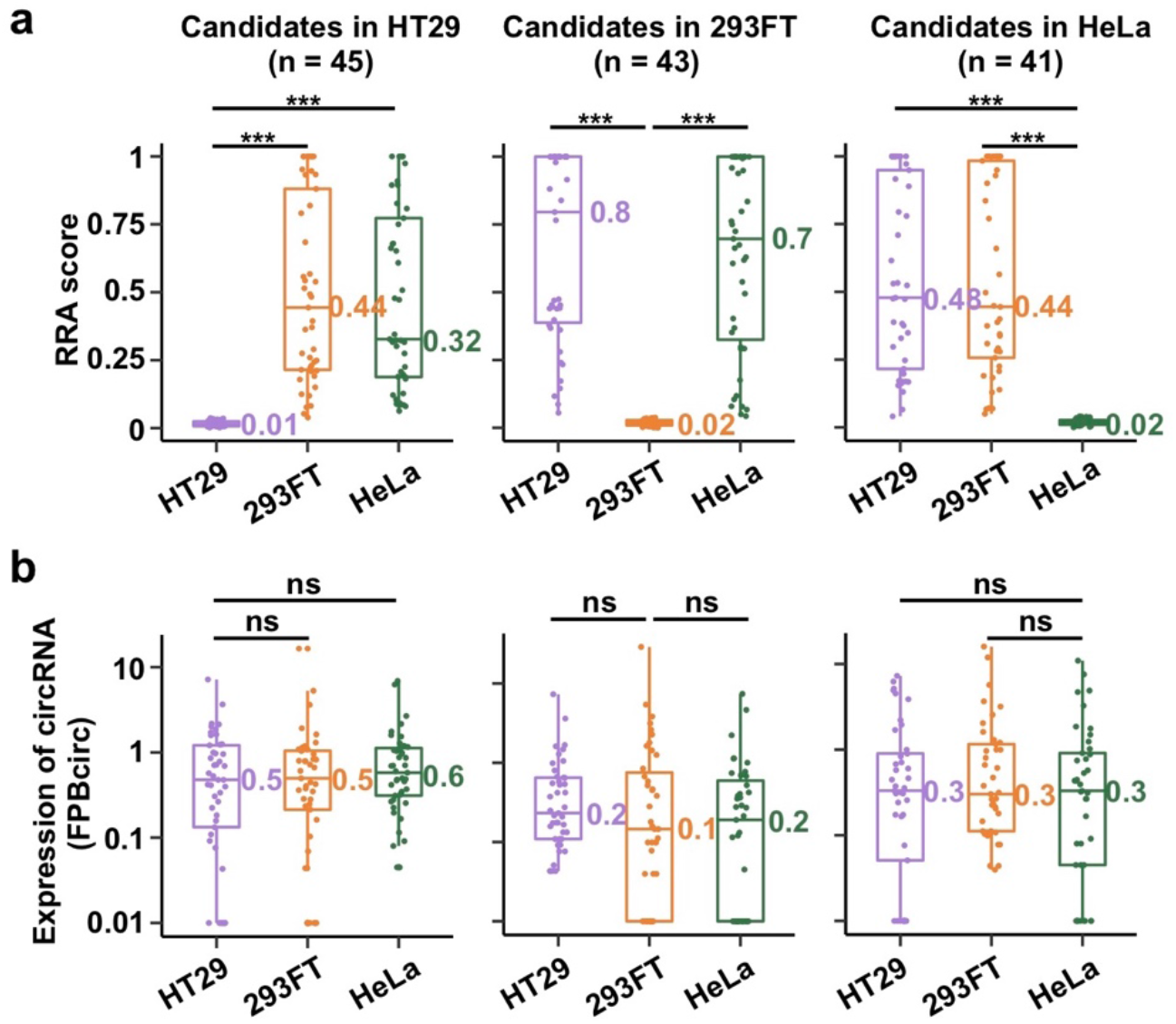
Distribution of RRA score of and expression level of circRNAs that have cell type-specific effects on cell growth. **a**, Distribution of the RRA score of circRNAs with cell-type specific effects in HT29 (45), 293FT (43) and HeLa (41) cell, respectively. **b**, Distribution of expression (shown by FPBcirc) of circRNAs shown in (**a**) in HT29, 293FT and HeLa cells.

**Extended Data Fig. 8.**
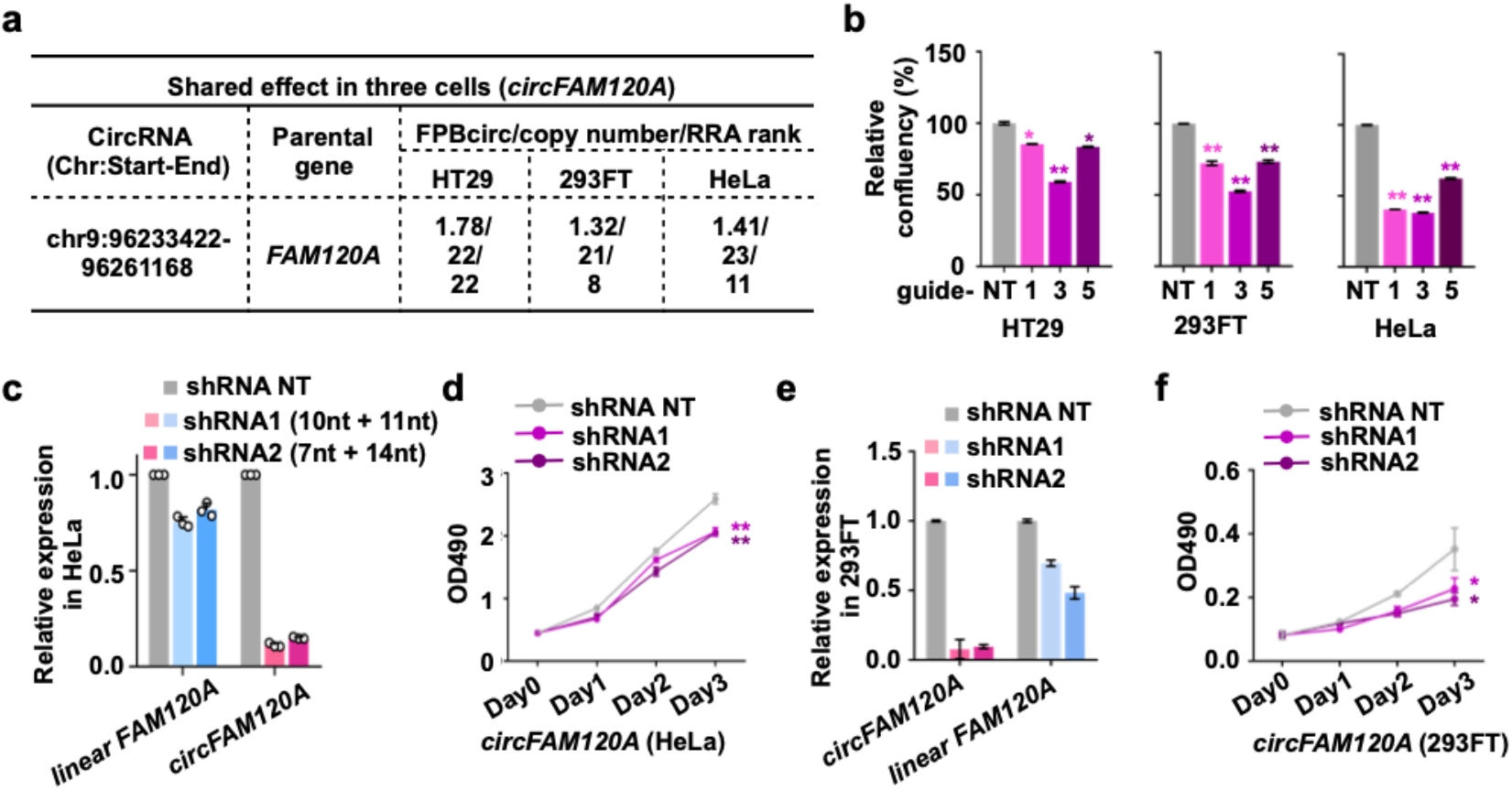
*circFAM120A* promotes cell proliferation. **a**, Summary of *circFAM120A* expression and growth defects in all three examined cell lines. **b**, Knockdown of *circFAM120A* by RfxCas13d/BSJ-gRNAs inhibited HT29, 293FT or HeLa cell proliferation, as revealed by cell confluency. **c**, Efficient knockdown of *circFAM120A* by shRNAs in HeLa cells. The expression of circRNAs and cognate linear RNAs was detected by qRT-PCR. **d**, Knockdown of *circFAM120A* by shRNAs inhibited cell proliferation in HeLa cells, as revealed by MTT assay. **e**, Efficient knockdown of *circFAM120A* by shRNAs in 293FT cells. The expression of circRNAs and cognate linear RNAs was detected by qRT-PCR. **f**, Knockdown of *circFAM120A* by shRNA inhibited cell proliferation in 293FT cells, as revealed by MTT assay. **b-d,f**, Values represent mean +/− SD with *n*=3, *: *P* < 0.05; **: *P* < 0.01. **c-e**, All transcript levels were normalized to *ACTB*, and values represent mean +/− SD with *n*=3.

**Extended Data Fig. 9.**
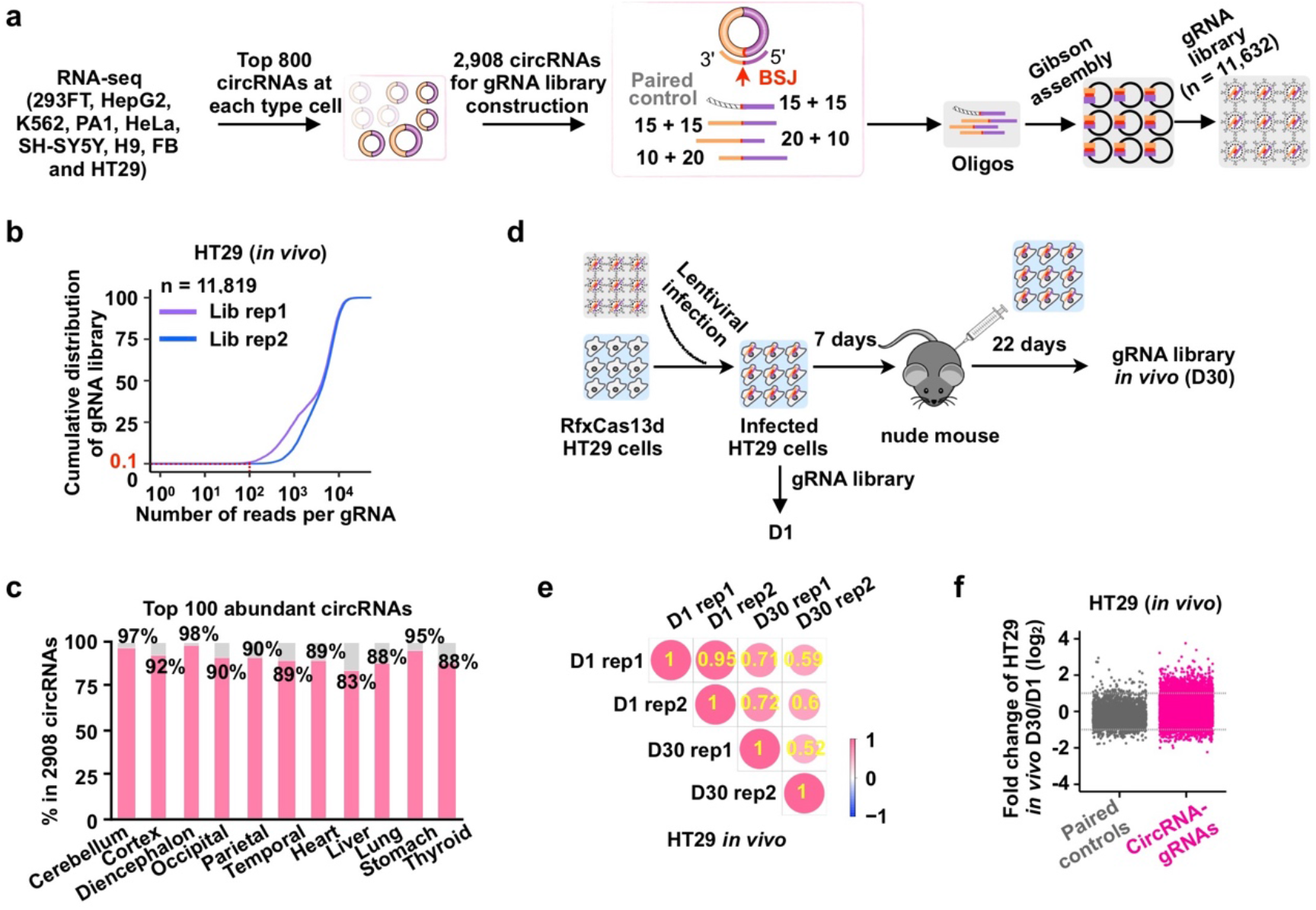
Overview of sequencing analyses of *in vivo* screens using BSJ-gRNA libraries targeting 2,908 circRNAs. **a**, Construction of an additional gRNA library targeting 2,908 circRNAs. One paired control gRNA (n=2,908) and three BSJ-gRNAs (circRNA gRNAs, n=8,724) were designed for each candidate circRNA. 100 negative control gRNAs were also included. **b**, Cumulative distribution of the number of reads per gRNA of constructed libraries. The red line indicates that less than 0.1% of gRNAs are covered by less than 100 reads. **c**, Representation of 2,908 candidate circRNAs in the library in different human tissues^43^. On average, over 90% of top 100 abundant circRNAs in each tissue were included in the list of 2,908 candidate circRNAs. **d**, *In vivo* screen of circRNAs important for cell growth and proliferation. The gRNA lentiviral library was individually delivered into HT29 cells stably expressing RfxCas13d. Infected cells were enriched after 7 days and injected subcutaneously to nude mouse for 22 days. Genomic DNAs from infected cells were extracted at day 1 (D1) and 30 (D30) for gRNA amplification and deep sequencing. **e**, The Pearson correlation coefficient (PCC) between replicates (rep) of D1 and D30 *in vivo* samples in HT29. Two biologically independent experiments were performed at D1 and D30. **f**, Scatter plot of fold change of paired controls and circRNA BSJ-gRNAs between D1 and D30 *in vivo* samples in HT29. The grey dashed lines indicate 2 or 0.5 fold change, respectively.

**Extended Data Fig. 10.**
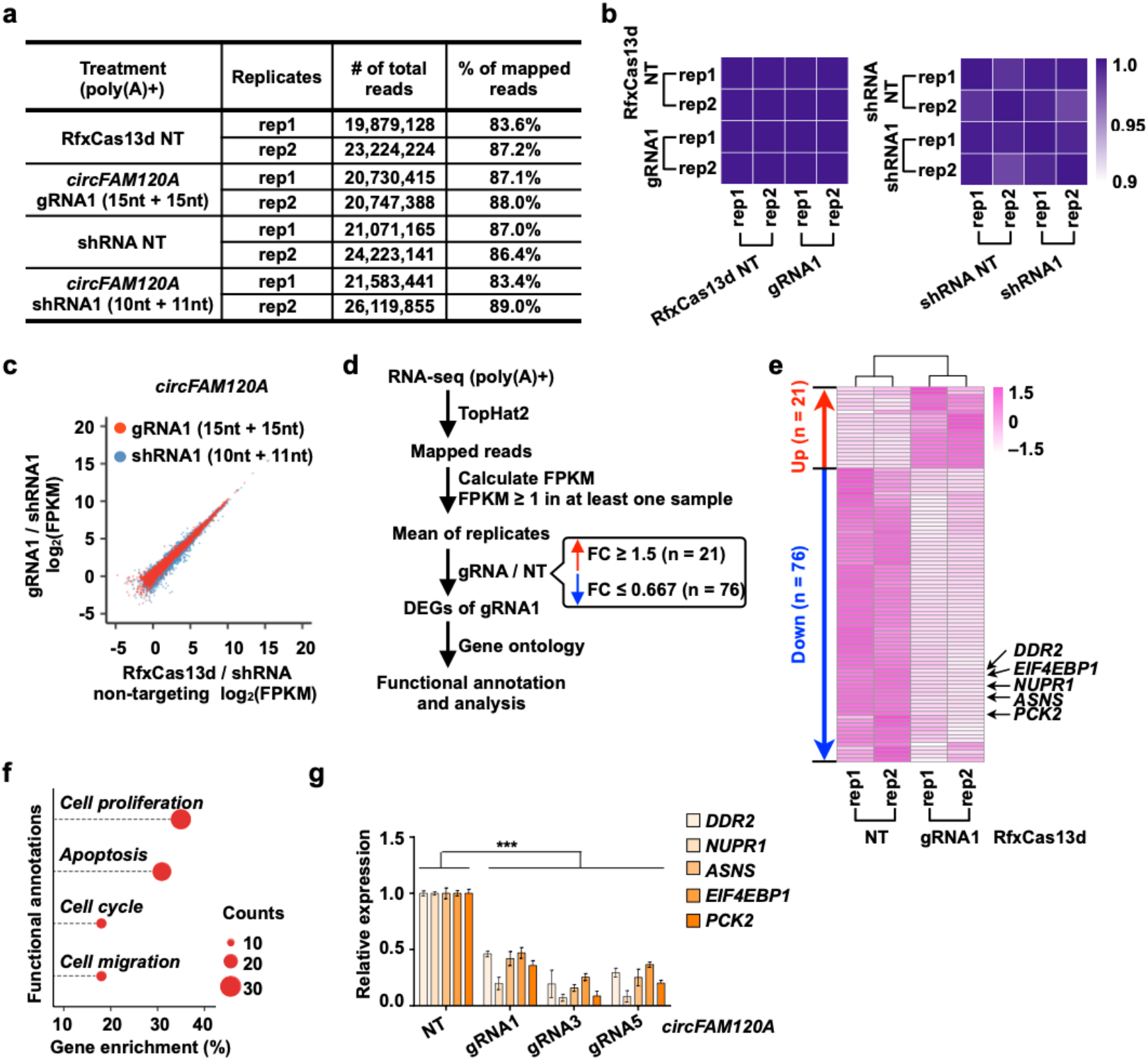
RNA-seq analysis of RfxCas13d/BSJ-gRNA- or shRNA-mediated *circFAM120A* knockdown. **a**, Mapping statistics of two biological replicates of the poly(A)+ RNA-seq datasets in 293FT cells with RfxCas13d/BSJ-gRNA- or shRNA-mediated *circFAM120A* knockdown. **b**, Heatmap of correlation (Kendall’s tau) for log_2_(FPKM) values of all linear mRNAs detected in RNA-seq libraries between targeting and non-targeting replicates for RfxCas13d/BSJ-gRNA- or shRNA-mediated *circFAM120A* knockdown. **c**, The expression levels in log_2_(FPKM) values of all genes detected in RNA-seq libraries of non-targeting control (x-axis) compared to *circFAM120A*-targeting conditions (y-axis) by RfxCas13d/BSJ-gRNA (red) or shRNA (blue). Means of two biological replicates were shown. **d**, A workflow shows the selection of candidate genes after *circFAM120A* knockdown by RfxCas13d/BSJ-gRNA. **e**, Heatmap of DEGs (n=97) detected in 293FT cells after knocking down *circFAM120A* targeted by RfxCas13d/BSJ-gRNA. **f**, Enrichment analysis of differentially expressed genes (DEGs) from RNA-seq after *circFAM120A* knockdown by RfxCas13d/BSJ-gRNA in 293FT cells. The x axis shows the ratio of the number of genes in a given category of functional annotations divided by the total number of DEGs. The y axis shows categories of functional annotations. **g**, Validation of DEGs associated with cell proliferation after *circFAM120A* knockdown by RfxCas13d/BSJ-gRNA in 293FT cells. All transcript levels were normalized to *ACTB*, and values represent mean +/− SD with *n*=3, ***: *P* < 0.001.

**Extended Data Fig. 11.**
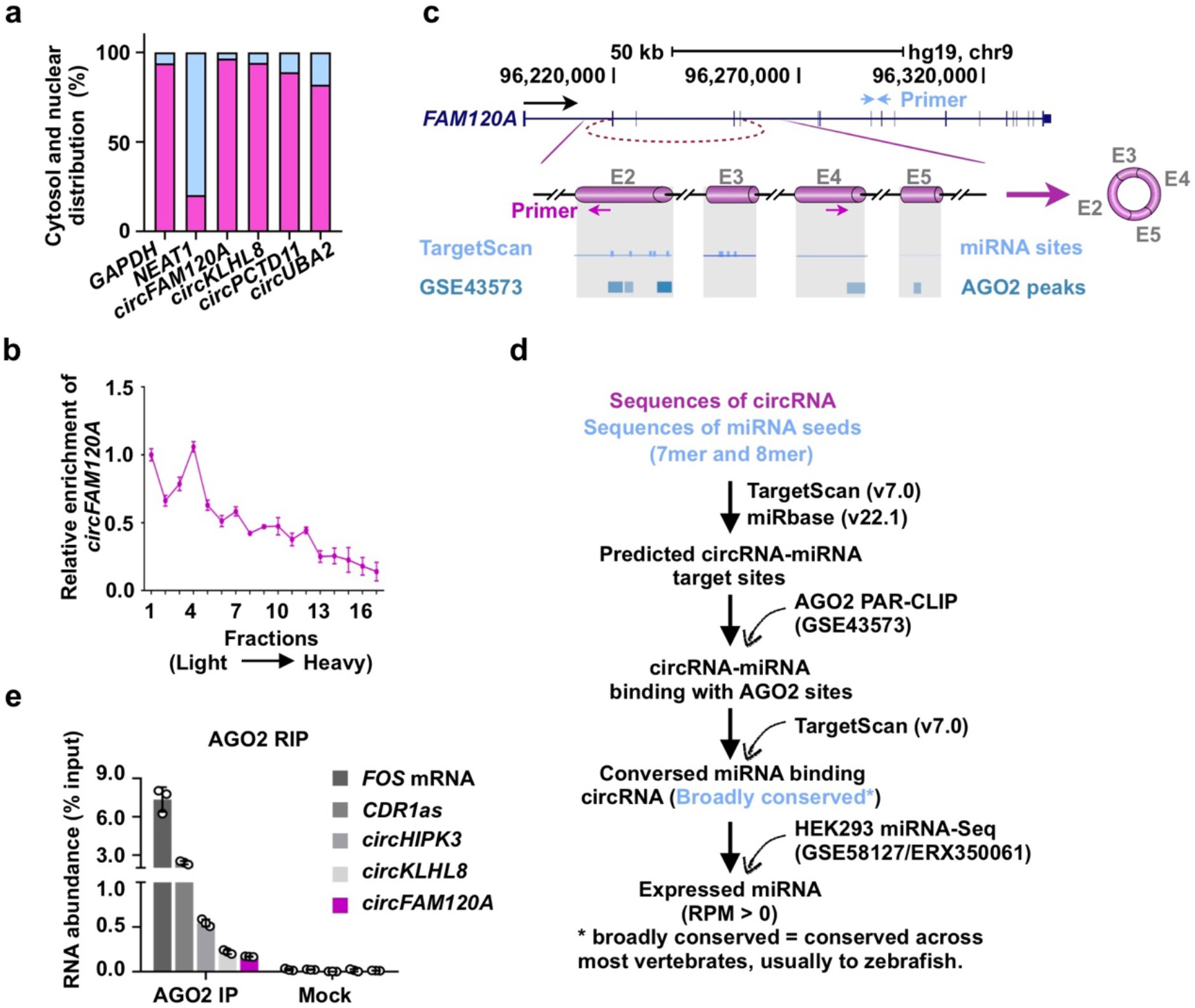
*CircFAM120A* does not act as miRNA sponge. **a**, Cytoplasmic distribution of *circFAM120A*. Total RNAs were separated into cytoplasmic and nuclear soluble fractions. The distribution of RNAs in the cytoplasmic and nuclear soluble fractions were detected by qRT-PCR. **b**, Cytoplasmic extracts from 293FT cells were loaded on 10%-45% sucrose gradients. The enrichment of *circFAM120A* in individual fractions were measured by qRT-PCR. Fraction density increases from left to right. **c**, Prediction of AGO2-binding peaks in *circFAM120A*. Top, genomics locus and diagram of linear *FAM120A* and *circFAM120A* (shown as cylinders in magenta). Blue and magenta arrows indicate location of primer for linear *FAM120A* or *circFAM120A*, respectively. Bottom, predicted miRNA target sites by TargetScan and AGO2 binding peaks from PAR-CLIP data in 293FT cells (GEO:GSE43573). **d**, A schematic drawing to show the strategy for predication between circular RNA and miRNA. Sequences of circRNA and miRNA seeds were used to predicted circRNA-miRNA target sites by TargetScan. The predicted binding sites were validated with AGO2 PAR-CLIP dataset (GSE43573), and broadly conserved miRNAs were used for the subsequent analysis. After that, expressed miRNAs were selected for future analysis in 293FT cells (GSE58127/ERX350061). **e**, *CircFAM120A* does not bind to AGO2 protein by RIP in 293FT cells using anti-AGO2 antibodies. The percentage of RIP-enriched RNAs relative to input was quantified by qRT-PCR.

**Extended Data Fig. 12.**
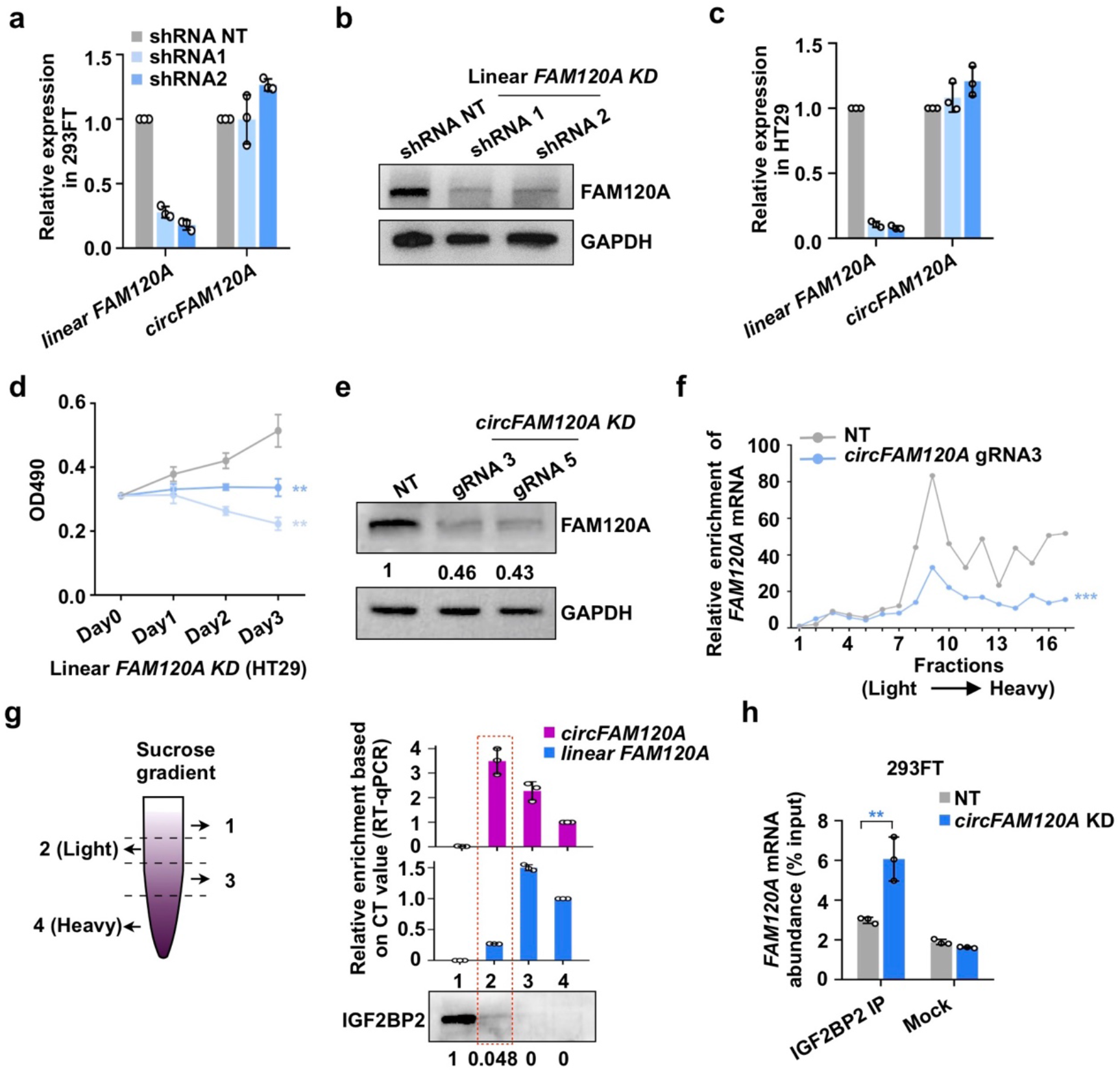
*CircFAM120A* promotes cell proliferation by regulating its parental gene translation in an IGF2BP2-dependent manner. **a**, Efficient knockdown of *FAM120A* by shRNAs in 293FT cells. The expression of circRNAs and cognate linear RNAs was detected by qRT-PCR, all transcript levels were normalized to *ACTB*, and values represent mean +/− SD with *n*=3. **b**, *FAM120A* knockdown by two different shRNAs in 293FT cells, revealed by WB. **c**, Efficient knockdown of *FAM120A* by shRNAs in HT29 cells. The expression of circRNAs and cognate linear RNAs was detected by qRT-PCR. **d**, Knockdown of *FAM120A* by shRNAs inhibited cell proliferation in HT29 cells, as revealed by MTT assay, and values represent mean +/− SD with *n*=3, **: *P* < 0.01. **e**, Knockdown of *circFAM120A* leads to reduced expression of FAM120A protein in HT29 cells. **f**, Knockdown of *circFAM120A* leads to reduced enrichment of linear *FAM120A* on polyribosomes. Cytoplasmic extracts from 293FT cells treated with NT-gRNA (gray) or *circFAM120A*-gRNA3 (blue), were loaded on 10%-45% sucrose gradients. Enrichments of *FAM120A* mRNA in individual fractions were measured by qRT-PCR. Fraction density increases from left to right. Statistically significance was assessed using Wilcoxon test with R platform (R v.3.5.1) and values represent mean +/− SD with *n*=3, ***: *P* < 0.001. **g**, The distribution of IGF2BP2 protein, *circFAM120* and *FAM120A* mRNA in four fractions of polysome profiling of 293FT cells. The schematic of the sucrose gradients used to segregate fractions in polysome profiling assay is shown on left. **h**, Knockdown of *circFAM120A* leads to increased interaction between cognate *FAM120A* mRNA and IGF2BP2 in 293FT cells. The abundance of *FAM120A* mRNA was measured by qRT-PCR, and values represent mean +/− SD with *n*=3, **: *P* < 0.01.

**Extended Data Fig. 13.**
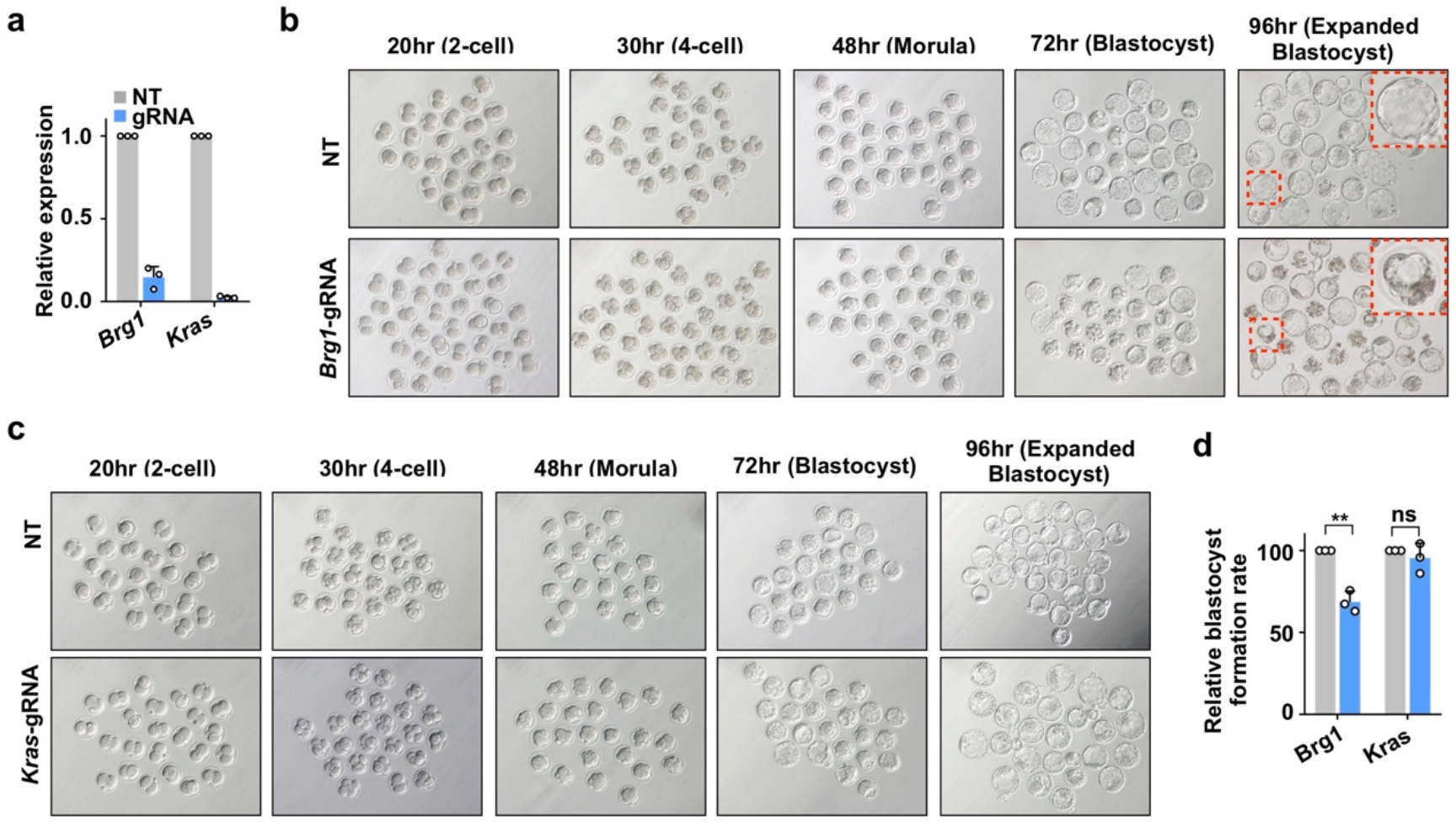
Application of RfxCas13d/gRNA to interfere linear RNA expression during mouse preimplantation development. **a**, Knockdown of *Brg1* and *Kras* by RfxCas13d/gRNA in zygotes. Expression levels of mRNAs were detected by qRT-PCR. All transcript levels are relative to *ACTB*, and values represent mean +/− SD with *n*=3. **b**, Representative images of reduced blastocyst formation 96h after microinjection of RfxCas13d mRNA and the gRNA targeting *Brg1* mRNA into mouse zygotes. An example of failed blastocyst formation is shown by red arrows and enlarged view. **c**, Representative images of embryogenesis after microinjection of RfxCas13d mRNA and the gRNA targeting *Kras* mRNA into mouse zygotes. Of note, no aberrant mouse preimplantation development was observed. **d**, Effect of *Brg1* and *Kras* knockdown on blastocyst formation 96h after microinjection of RfxCas13d mRNA and BSJ-gRNAs into mouse zygotes.

**Extended Data Fig. 14.**
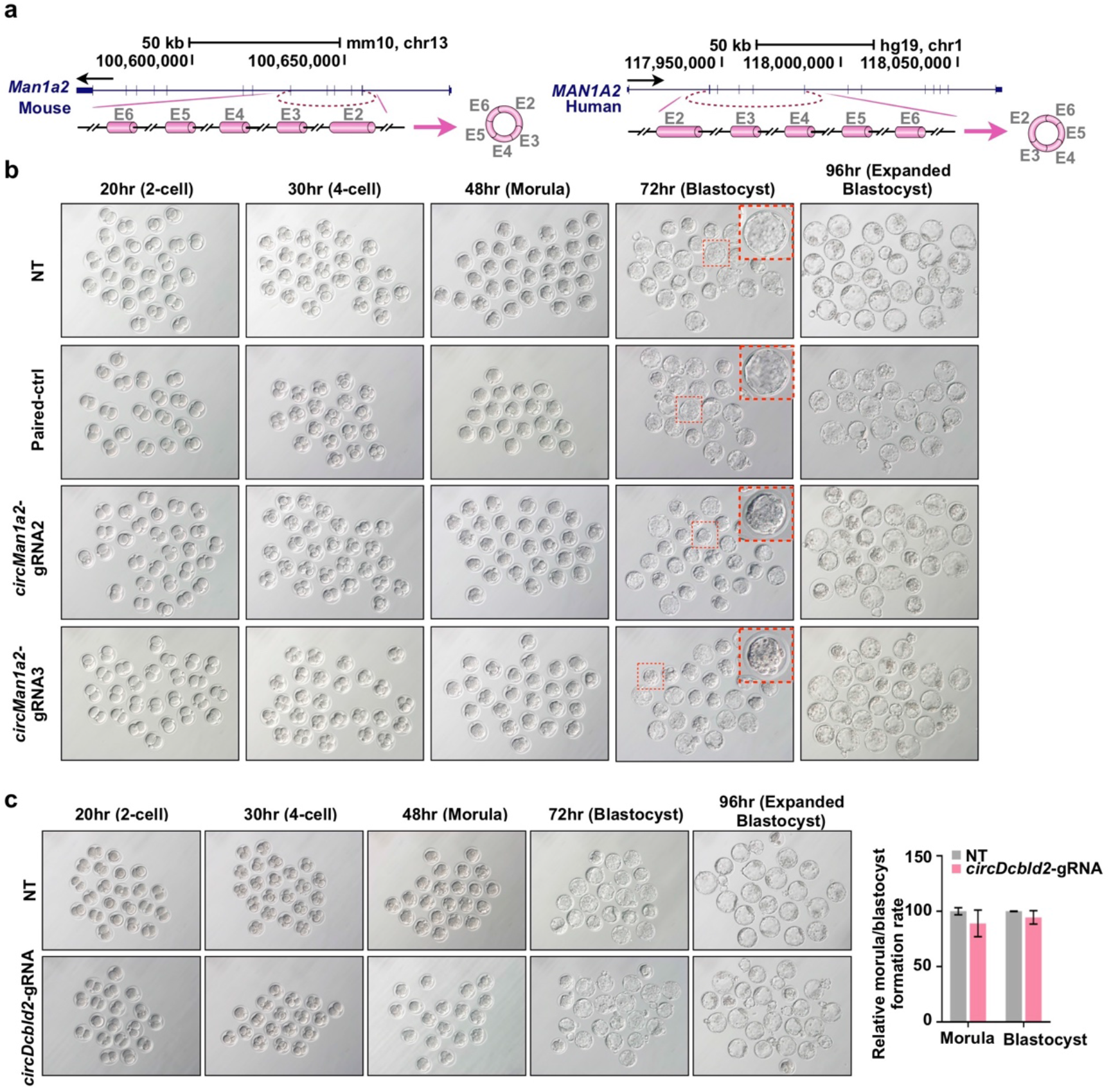
Application of RfxCas13d/gRNA to interfere circRNA expression during mouse preimplantation development. **a**, Genomic loci and diagrams of mouse *circMan1a2* and human *circMAN1A2* were shown as cylinders in magenta. **b**, Knockdown of *circMan1a2* in zygotes led to reduced blastocyst formation 72h after microinjection of RfxCas13d mRNA and the corresponding BSJ-gRNAs into mouse zygotes. Representative images of at 2-cell, 4-cell, morula and blastocyst stages are shown; an example under each condition is highlighted by red line and enlarged view. **c**, Images show the normal embryonic morphologies at 2-cell, 4-cell, morula and blastocyst stages under *circDcbld2* knockdown by RfxCas13d/BSJ-gRNA in zygotes. The effect of *circDcbld2* knockdown on morula and blastocyst formation was showed on right.

## Supplementary Tables and Table Legends

**Supplementary Table 1. List of sequences of RfxCas13d/BSJ-gRNA library for 762 circRNAs in this study**.

**a**, The expression (FPBcirc) of 762 circRNAs in HT29, 293FT and HeLa cells.

**b**, The sequences of RfxCas13d/BSJ-gRNA library for 762 circRNAs.

**Supplementary Table 2. MAGeCK results of negatively selected circRNAs in HT29, 293FT and HeLa cell lines.**

**a**, The robust rank aggregation (RRA) scores of ranked circRNAs in HT29 cells (D30/D1).

**b**, RRA scores of ranked circRNAs in 293FT cells (D30/D1).

**c**, RRA scores of ranked circRNAs in HeLa cells (D30/D1).

**Supplementary Table 3. List of the normalized reads of top 50 candidate circRNAs identified by RfxCas13d/BSJ-gRNA screening in HT29, 293FT or HeLa cell lines.**

**a**, Normalized reads of gRNAs for 50 candidate circRNAs in HT29 cells.

**b**, Normalized reads of gRNAs for 50 candidate circRNAs in 293FT cells.

**c**, Normalized reads of gRNAs for 50 candidate circRNAs in HeLa cells.

**Supplementary Table 4.** List of validated circRNA candidates by RfxCas13d/BSJ-gRNA in HT29, HeLa and 293FT cells. Their parental genes, genomic locations, FPBcirc, RRA ranks are listed. Most of examined candidate circRNAs can be validated by RfxCas13d with at least one gRNA in HT29, HeLa and 293FT cells, respectively.

| Parental gene | CircRNA<br>(Chr:Start-End)<br>Conserved (mm10) | # gRNAs with<br>KD efficiency (>50%)<br>/# total validated gRNAs | Inhibition of<br>cell<br>proliferation | FPBcirc |  |  | RRA rank<br>*In vitro* L I |
| --- | --- | --- | --- | --- | --- | --- | --- |
|  |  |  |  | HT29 | 293FT | HeLa |  |
| TJP1 | chr15:30053341-30092905 | 1/1 gRNA | ✓ | 0.41 | — | 0.71 | 6 |
| NFATC3 | chr16:68155889-68157024 | 1/1 gRNA | ✗ | 1.53 | 1.58 | 0.78 | 10 |
| DCBLD2 | chr3:98568304-98600611 | 1/1 gRNA | ✓ | 0.98 | 0.41 | — | 15 |
| CSNK1G3 | chr5:122881110-122911657 | 1/1 gRNA | ✓ | 0.68 | 0.10 | 0.28 | 17 |
| ERBB2IP | chr5:65284462-65290692 | 1/1 gRNA | ✓ | 2.15 | 0.33 | 1.91 | 20 |
| FAM120A | chr9:96233422-96261168 | 3/3 gRNA | ✓ | 1.78 | 1.32 | 1.41 | 22 |
| AMD1 | chr6:111208707-111211559 | 1/1 gRNA | ✓ | 1.03 | 0.95 | 0.78 | 27 |
| BPTF | chr17:65941524-65944422 | 1/1 gRNA | ✓ | 0.31 | 0.08 | 0.15 | 30 |
| PCMTD1 | chr8:52773404-52773806 | 1/1 gRNA | ✗ | 2.67 | 2.12 | 5.28 | 48 |
| KLHL8 | chr4:88116475-88116842 | 3/3 gRNA | ✓ | 0.44 | 1.52 | 1.47 | 1 |
| AMBRA1 | chr11:46515157-46529920 | 0/1 gRNA | ✗ | 0.28 | 0.36 | 0.37 | 4 |
| FAM120A | chr9:96233422-96261168 | 3/3 gRNA | ✓ | 1.78 | 1.32 | 1.41 | 8 |
| FAF1 | chr1:51253671-51267349 | 1/1 gRNA | ✓ | — | 1.60* | 0.09 | 10 |
| PMS1 | chr2:190656515-190682906 | 1/1 gRNA | ✓ | 1.12 | 1.04 | 1.78 | 12 |
| ATXN2 | chr12:111990083-111993723 | 1/1 gRNA | ✓ | 2.21 | 0.65 | 3.13 | 16 |
| UBAP2 | chr9:33971648-33973235 | 1/1 gRNA | ✓ | 0.20 | 4.66 | 0.44 | 17 |
| PDCD11 | chr10:105197771-105198565 | 1/1 gRNA | ✓ | — | 0.17 | 0.09 | 21 |
| EXOC6B | chr2:72945231-72960247 | 1/1 gRNA | ✓ | 0.73 | 1.66 | 2.60 | 28 |
| CCNB1 | chr5:68470703-68471364 | 1/1 gRNA | ✓ | 0.99 | 1.47 | 1.62 | 37 |
| CLASP1 | chr2:122363276-122363756 | 1/1 gRNA | ✓ | 0.42 | 0.57 | 0.46 | 9 |
| FAM120A | chr9:96233422-96261168 | 3/3 gRNA | ✓ | 1.78 | 1.32 | 1.41 | 12 |
| XPO1 | chr2:61749745-61761038 | 1/1 gRNA | ✓ | 4.69 | 4.52 | 3.65 | 13 |
| KLHL8 | chr4:88116475-88116842 | 3/3 gRNA | ✓ | 0.44 | 1.52 | 1.47 | 20 |
| ATXN10 | chr22:46085591-46098727 | 0/1 gRNA | ✗ | 0.31 | 0.44 | 0.79 | 22 |
| ASPH | chr8:62593526-62596747 | 1/1 gRNA | ✓ | 4.88 | 1.95 | 11.93 | 29 |
| HIPK3 | chr11:33307958-33309057 | 3/3 gRNA | ✓ | 10.92 | 6.23 | 15.97 | 35 |
| MIB1 | chr18:19345732-19359646 | 1/1 gRNA | ✓ | 1.75 | 7.26 | 3.18 | 36 |
| CRKL | chr22:21288066-21288532 | 1/1 gRNA | ✓ | 3.25 | 4.93 | 1.20 | 46 |

**Supplementary Table 5. List of sequences of RfxCas13d/BSJ-gRNA library for 2908 circRNAs in this study.**

**a**, The sequences of RfxCas13d/BSJ-gRNA library for 2908 circRNAs.

**b**, The robust rank aggregation (RRA) scores of ranked circRNAs in HT29 *in vivo* (D30/D1).

**c**, The top 200 robust rank aggregation (RRA) scores of ranked circRNAs in HT29 *in vivo* (D30/D1).

**d**, Overlap of top 50 *in vitro* and top 200 *in vivo* robust rank aggregation (RRA) scores of ranked circRNAs in HT29 (D30/D1).

**Supplementary Table 6.**
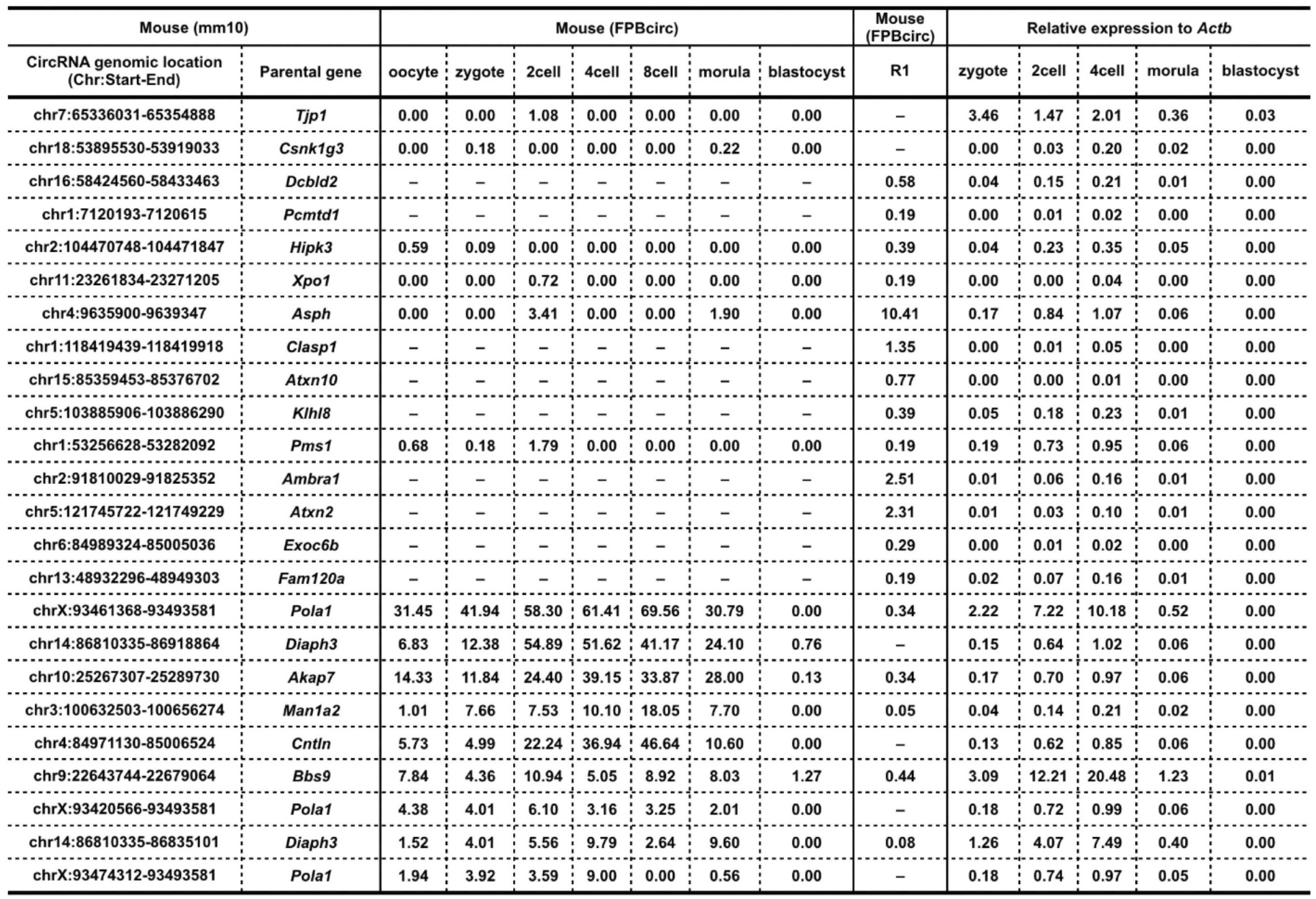
Expression of 24 circRNA candidates in mouse preimplantation development.

**Supplementary Table 7.**
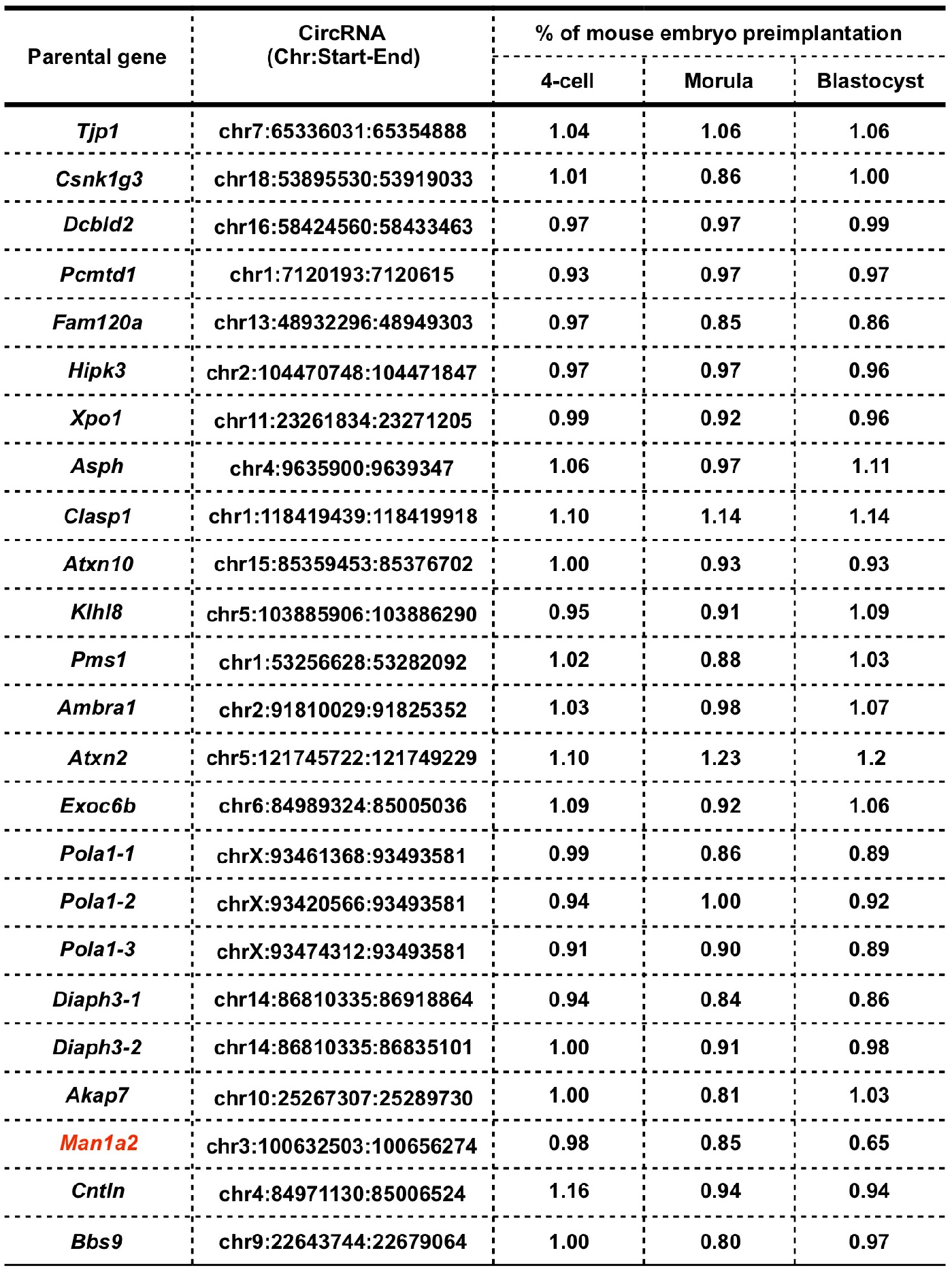
Screening of circRNAs with potential regulatory roles in mouse embryo preimplantation development by RfxCas13d/BSJ-gRNA KD each of 24 circRNA candidates in mouse zygotes.

**Supplementary Table 8. Expression of linear genes upon *circFAM120A* knockdown by RfxCas13d or shRNA in 293FT cells.**

**a**, List of linear genes after *circFAM120A* knockdown in RfxCas13d NT and gRNA1.

**b**, List of linear genes after *circFAM120A* knockdown in shRNA NT and shRNA1.

**Supplementary Table 9. List of sequences used in this study.**

**a**, gRNA sequences.

**b**, shRNA sequences.

**c**, Primers for qRT-PCR and NB probes.

